# Prognosis prediction using autophagy gene expression in osteosarcoma

**DOI:** 10.1101/2025.11.05.685081

**Authors:** Qingzhu Liu, Cong Zhou, Ke Xu, Qikui Zhu, Junqin Lu, Chun Zhang, Wukun Xie, Guojiu Fang, Xue Zhou Chen, Dasheng Tian, Juehua Jing, Yize Li, Hongsheng Wang, Wenxiu Duan, Yihui Bi

## Abstract

Defining the transition from localized to metastatic osteosarcoma (OS) before overt dissemination is fundamental for improving survival. However, effective early diagnostic tools remain scarce, largely due to limited exploitation of the pre-metastatic tumor microenvironment’s own record of molecular barcode exposures encoded in cell-intrinsic stress states. We analyzed autophagy-dependent transcriptional datasets from 139 individuals to develop Auto-RS, a computational classifier that can integrate age and sex to deliver individualized risk management. The prognosis-interpretable features of Auto-RS recapitulate established molecular trajectories of metastasis at the single-cell level, capturing tumor cells’ intrinsic shift from proliferative to invasive states and revealing cooperative programs among cancer-associated fibroblasts and immune cells. More significantly, Auto-RS can expose chemotherapy vulnerabilities of newer drugs, providing a framework to prioritize therapeutics without direct testing in children. This framework brings a potential inflection point, where metastasis prediction and therapeutic stratification may converge to improve OS outcomes.

## Introduction

Osteosarcoma (OS) is rare but remains the most common primary malignant bone tumor, with an annual incidence of 0.2-3 cases per 100,000 children and 0.8-11 cases per 100,000 adolescents, which coincides with periods of rapid skeletal growth^1,2^. In the United States, an estimated 800-900 new cases are diagnosed annually, most in children and young adults. Current guidelines recommend multi-agent chemotherapy combined with complete surgical resection has increased 5-year survival for localized disease to ∼ 60%^3^. However, despite decades of therapeutic exploration, survival for patients with metastatic or recurrent OS has stagnated at < 20%^4^.

Most OS patients harbor clinically undetectable micro-metastases at presentation, a hallmark attributed to the tumor’s aggressive biology and profoundly unstable genomic architecture^5–8^. Recent consensus suggests that transformative advances in treatment are unlikely to result from intensification of antineoplastic chemotherapy, as the rarity of OS worldwide precludes large-scale trials of newer drugs, and companies prefer to avoid direct testing in children^9,10^. Instead, the post-genomic era is driving the development of stratification systems that categorize patients based on prognostic or biological features of OS, with the potential to impact outcomes^1,2,10,11^.

Yet, existing prognostic models for OS metastasis remain limited and suboptimal because they lacked systematic screening and in-depth interpretability^12^.

Here, we sought to establish an effective tool for early prediction of OS prognosis and to functionally map its biological relevance at the single-cell level within the tumor microenvironment.

## Results

### OS patient cohorts

We utilized ongoing Therapeutically Applicable Research To Generate Effective Treatments (TARGET) study information, curated by the U.S. National Cancer Institute (NCI), which comprises genomic profiles of 88 clinically annotated patients. In parallel, we incorporated whole-genome expression data from 53 high-grade OS cases provided by the EuroBoNet consortium and GEO database (GSE21257)^13^. Two TARGET cases were excluded due to missing survival data. Baseline clinicopathological characteristics of both cohorts are summarized in ***Table 1***.

**Table 1:**
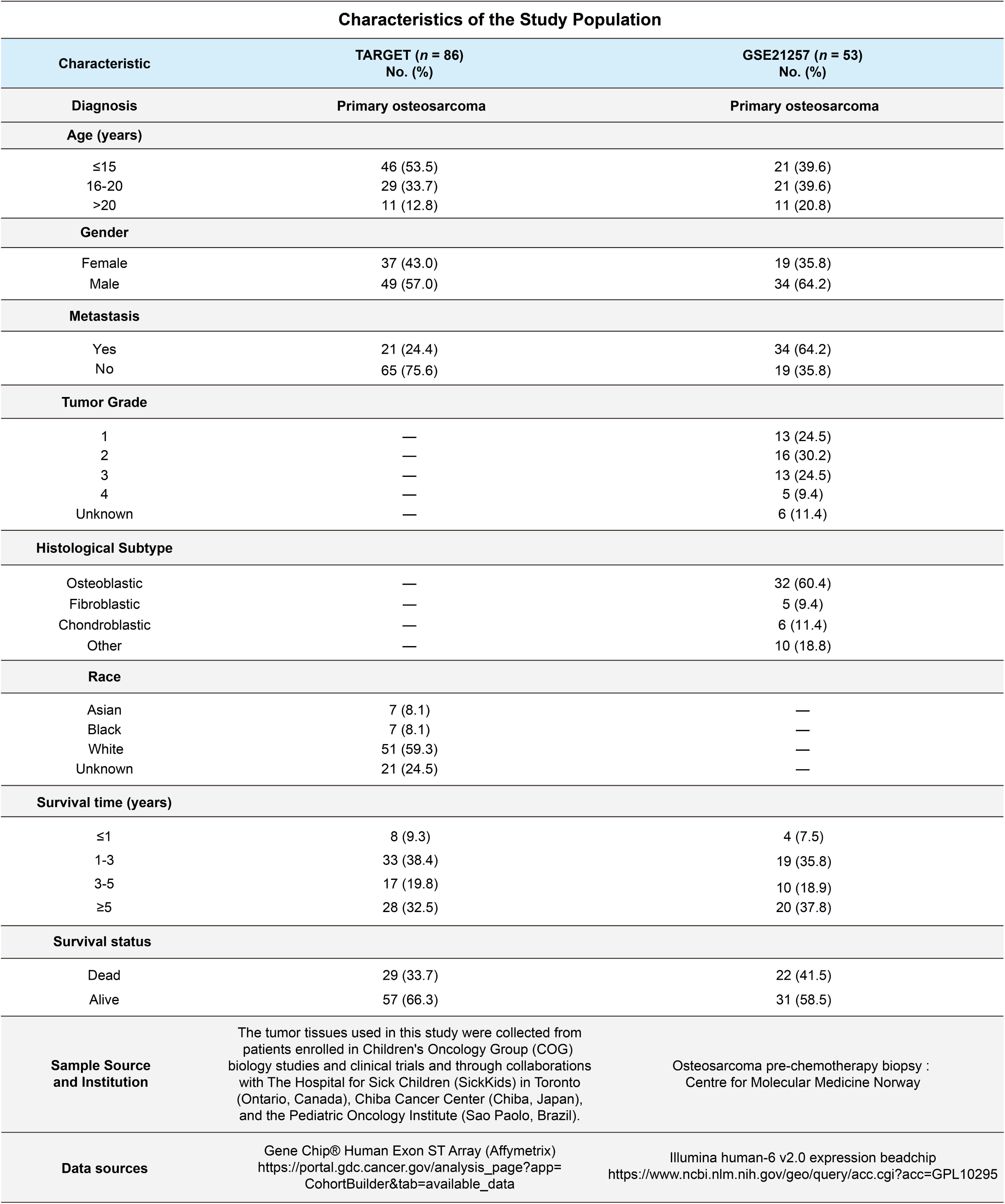
Clinical and pathological characteristics of patients and their tumors.

### Autophagy gene dysregulation characterizes metastatic OS

Patients with metastasis had significantly shorter survival than those without (TARGET, *P* = 0.0018, GSE21257, *P* = 8.4e-07; Fig. S1A and S1B). Multivariable Cox regression confirmed metastasis as an independent predictor of poor prognosis (*P* < 0.001), whereas age and sex were not statistically significant (Gender*, P* = 0.574, Age*, P* = 0.976; Fig. S1C). On this basis, we developed a basic prognostic model (BpM) as a nomogram to estimate 1-, 3-, and 5-year overall survival (Fig. S1D). Although age and sex did not independently affect outcome, both were retained given prior evidence of clinical relevance in OS, thereby enhancing real-world applicability^2^. Metastatic status carried the greatest prognostic weight, producing the widest spread of total scores, with survival probability declining steeply as scores increased (***Table S1***). Model calibration using 1,000 *Bootstrapping* showed close agreement between predicted and observed outcomes at higher risk levels (survival probability 0.0-0.6), with deviations emerging at the opposite end of the risk spectrum (survival probability 0.6-1.0) (Fig. S1E).

We next profiled the transcriptional differences between metastatic and non-metastatic cases to identify molecular features that could refine the BpM, particularly in cases lacking explicit metastasis annotation, enabling more individualized risk stratification. Our earlier attempt to derive model genes from global transcriptomic data was abandoned due to limited interpretability, as discussed later (see ***Discussion***). Given the well-established role of autophagy program in metastatic progression across cancers^14–21^, we focused on the dysregulation of autophagy genes in OS. To this end, expression data were integrated, with batch effects corrected using the *BatchServer* and the *sva* (Fig. S1F and S1G). Cross-referencing with the Human Autophagy Database yielded 193 autophagy genes (Fig. S1H, ***Table S2*** and ***S3***). Differentially expressed gene (DEG) analysis (*limma*) identified 12 upregulated and 17 downregulated autophagy genes in metastatic versus non-metastatic cases (Fig. S1I). We further visualized the gene-wise expression trajectories stratified by metastasis status, and examined correlations of DEGs with sex (gender) and survival status (Fustat) (Fig. S1J). These analyses underscored both the sensitivity of autophagy gene expression to metastatic progression and considerable heterogeneity among patients.

### Screening prognostic signatures-associated autophagy genes

To derive a parsimonious yet robust prognostic signature from autophagy genes, we developed a multistep feature selection framework that integrated least absolute shrinkage and selection operator (LASSO) regression with multivariate Cox proportional hazards modeling. This enabled simultaneous dimensionality reduction and capture of survival-associated signals. The workflow proceeds in three stages: (i) random partitioning of the integrated cohorts into training and testing subsets (Fig. 1A and ***Methods***, ***Table S4***); (ii) identification of prognosis-associated autophagy genes and construction of a risk score (*Auto-RS*) formula (Fig. 1B-E); and (iii) evaluation of predictive performance in both subsets (Fig. 1F-I). During feature selection, LASSO regression was employed to mitigate overfitting and to address multicollinearity inherent in high-dimensional transcriptomic data (Fig. 1C). By imposing a regularization penalty, LASSO shrinks the coefficients of less informative variables toward zero, yielding a sparse set of candidate predictors. The analysis was conducted using the *glmnet*, with 10-fold cross-validation to determine the optimal regularization parameter (λ). Genes with non-zero coefficients at λ_min (the value minimizing the cross-validated error) were retained (Fig. 1D). To further improve interpretability and reduce complexity, LASSO-selected features were subjected to bidirectional stepwise multivariate Cox regression, guided by minimization of the Akaike Information Criterion (AIC) (Fig. 1E). This refinement step prioritized the most informative predictors while penalizing excessive model complexity.

**Fig. 1:**
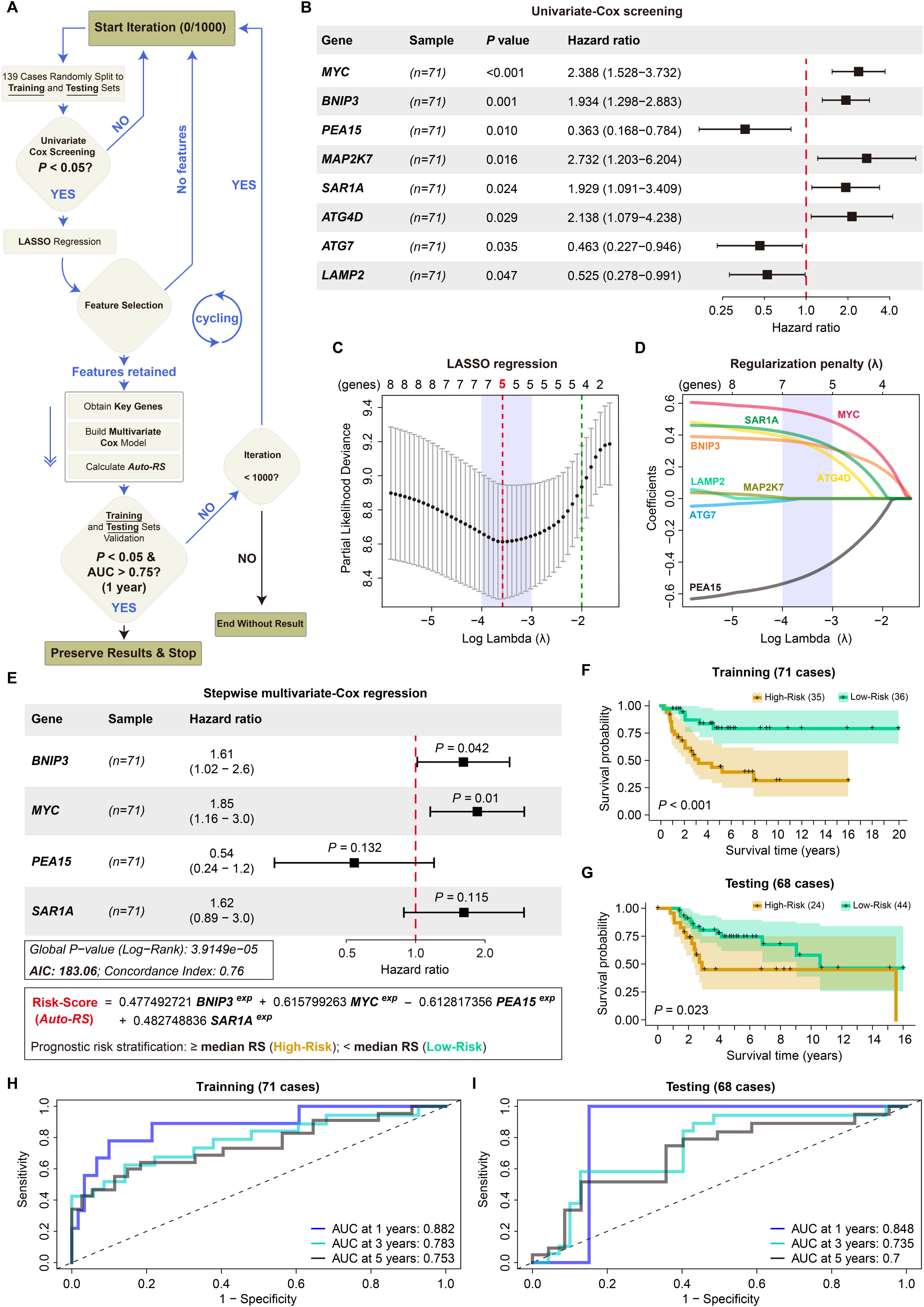
Self-supervised machine learning identifies prognostic autophagy genes. **A**, Workflow of automated screening integrating LASSO and Cox modeling. Rectangles denote operational steps, diamonds indicate decision nodes, and arrows mark flow direction (detailed in ***Methods***) **B**, Forest plot of eight autophagy genes significantly associated with OS survival by univariate Cox analysis (Hazard ratio with 95% CI, Wald test) **C**, Tenfold cross-validation for the LASSO regression model. The partial likelihood deviance curve is plotted against log(λ), dotted vertical lines represent the values of λ corresponding to the minimum deviance (red) and the one-standard-error criterion (green). **D**, LASSO coefficient profiles of candidate genes. Each colored line represents one gene; coefficients shrink toward zero with increasing λ, retaining only the most predictive genes. **E**, Multivariate Cox regression identifying four independent prognostic genes (*BNIP3*, *MYC*, *PEA15*, *SAR1A*) with corresponding Hazard ratios (95% CI, Wald test). The *Auto-RS* was calculated as the sum of gene expression weighted by Cox coefficients, dichotomized by median into high- vs. low-risk groups. The global *P* value was obtained by the log-rank test. **F**, **G**, Kaplan–Meier survival curves of training set (*n* = 71) and testing set (*n* = 68), showing significantly worse survival in the high-risk group (log-rank *P* < 0.05). **H**, **I**, Time-dependent ROC curves assessing model discrimination at 1-, 3-, and 5-year survival. AUC values demonstrate high predictive accuracy in both training and testing cohorts.

This framework screened 20 candidate gene sets (***Table S5***), and those can stratify survival (*P* < 0.05) were retained (***Table S6***). Among these, models with ΔAIC ≤ 10 were further considered, and the one with the highest concordance index (C-index) was ultimately selected (***Table S7***). Univariate Cox analysis identified 8 survival-associated autophagy genes, and LASSO regression reduced these to 5 candidates (Fig. 1B-D). Stepwise multivariate Cox regression further refined the model, resulting in a four-gene signature comprising *BNIP3*, *MYC*, *PEA15*, and *SAR1A* (Fig. 1E). Among them, *PEA15* (Hazard ratio = 0.54) acted as a protective factor, while *BNIP3* (Hazard ratio = 1.61), *MYC* (Hazard ratio = 1.85), and *SAR1A* (Hazard ratio = 1.62) were risk factors. The *Auto-RS,* calculated by weighting gene expression with Cox coefficients (β), effectively stratified patients in both training and testing cohorts (*P* < 0.05; Fig. 1F and 1G), with strong predictive accuracy (training AUCs: 0.882, 0.783, 0.753; testing AUCs: 0.848, 0.735, 0.700; Fig. 1H and 1I). Notably, *Auto-RS* reclassified 19 metastatic patients as low-risk and 33 non-metastatic patients as high-risk, indicating its ability to capture heterogeneity beyond clinical metastasis (Fig. S2A and S2B, ***Table S8***). Kaplan–Meier and time-dependent ROC analyses further confirmed its robust predictive performance in both the integrated cohort (AUCs: 0.869, 0.742, 0.701; Fig. S2C and S2D) and the individual group (Fig. S2E-H).

In summary, *BNIP3*, *MYC*, *PEA15*, and *SAR1A* constitute a four-gene prognostic signature that refines risk stratification and captures molecular heterogeneity in OS.

### *Auto-RS* is an independent prognostic factor

To assess clinical utility, we incorporated *Auto-RS* into prognostic modeling for OS. In univariate analysis, *Auto-RS* showed significant predictive power (Hazard ratio = 1.091, 95% CI: 1.047– 1.136, *P* < 0.001) (Fig. 2A). Multivariate Cox regression confirmed that both metastasis (Hazard ratio = 5.340, 95% CI: 2.840–10.042, *P* < 0.001) and *Auto-RS* (Hazard ratio = 1.058, 95% CI: 1.012–1.107, *P* < 0.05) were independent predictors of survival (Fig. 2B). Time-dependent ROC analysis further identified *Auto-RS* as the most accurate single predictor (AUC = 0.88), outperforming metastasis (0.83), gender (0.45), and age (0.39) (Fig. 2C). We next integrated *Auto-RS* into a complete prediction model (CpM), generating individualized nomograms for 1-, 3-, and 5-year survival (Fig. 2D) and total point scores for all 139 patients (***Table S1***). Calibration analysis demonstrated strong agreement between predicted and observed survival, correcting deviations observed in BpM (survival probability: 0.6-1.0) (Fig. 2E). Notably, CpM also produced a wider spread of predicted survival probabilities, reflecting improved capture of inter-patient heterogeneity.

**Fig. 2:**
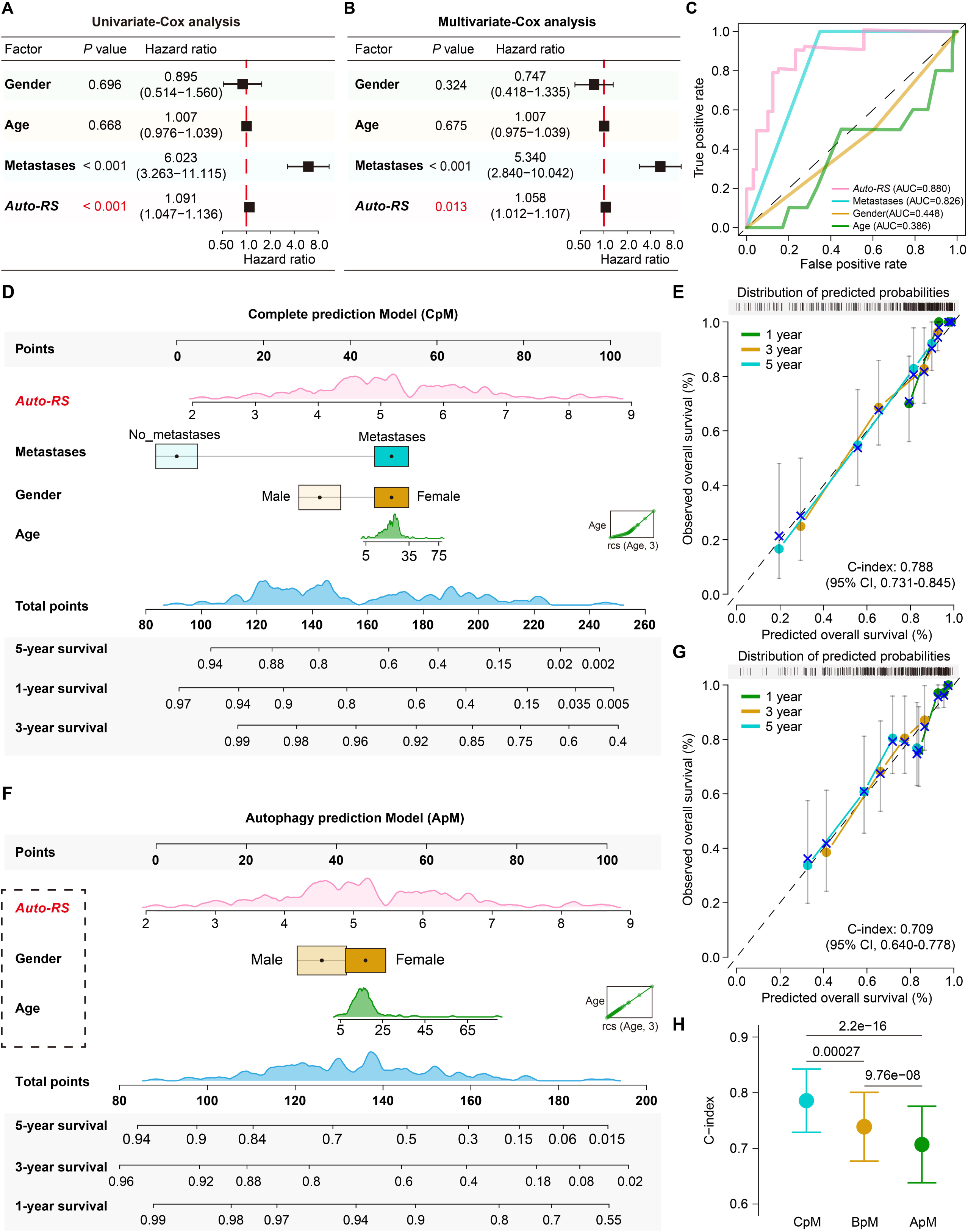
*Auto-RS* capture heterogeneity beyond clinical metastasis. **A**, Distribution of *Auto-RS* and survival status for all 139 OS patients. Low- and high-risk groups are shown in green and yellow, respectively; alive and deceased patients are shown in pink and black. **B**, Heatmap of the four *Auto-RS* component genes, row-wise Z-score normalized across samples and grouped by *Auto-RS*. Gender, metastatic status, and survival status (Fustat) are indicated above the heatmap. **C**,**E**,**G**, Kaplan–Meier survival curves for the combined cohort (*n* = 139), TARGET cohort (*n* = 86), and GSE21257 cohort (*n* = 53). High-risk patients exhibited significantly shorter survival compared with low-risk patients. *P*-values were calculated using the log-rank test; *P* < 0.05 was considered significant. **D**,**F**,**H**, Time-dependent ROC curves for the three cohorts. Blue, cyan, and light gray lines represent 1-, 3-, and 5-year survival predictions, respectively. AUC values reflect model performance, with higher values indicating better predictive accuracy.

Recognizing that clinical metastasis diagnosis is often delayed and survival improvements for metastatic OS have remained stagnant, we developed the autophagy prediction Model (ApM), which combines *Auto-RS* with age and gender while deliberately excluding metastasis to enable prognostic assessment. ApM retained strong predictive power with a broad range of individualized survival estimates, demonstrating its ability to capture prognostic heterogeneity independently of metastasis (Fig. 2F and 2G). Although ApM’s C-index (0.709; 95% CI: 0.640-0.778) was slightly lower than that of BpM (0.741; 95% CI: 0.679-0.803) and CpM (0.788; 95% CI: 0.731-0.845), ApM still achieved reliable risk stratification (Fig. 2H). Patient-level comparisons across the three models are provided in ***Table S1*.**

In summary, *Auto-RS* confers independent prognostic value, while ApM demonstrates the feasibility of metastasis-independent prediction, offering a framework for potentially early risk stratification.

### Cellular origin of *Auto-RS* signals

To determine what cells mainly deploy the *Auto-RS*, we carried out single-cell RNA-seq (scRNA-seq) analysis on six primary OS tumors^22^. Unsupervised clustering identified eight distinct populations from 27,719 cells, including myeloid cells (49.53%), osteoblastic OS cells (22.33%), NK/T cells (17.92%), cancer-associated fibroblasts (CAFs, 3.3%), plasma cells (2.54%), osteoclastic cells (1.9%), B cells (1.65%), and endothelial cells (0.82%) (Fig. 3A and S3A-C). *Auto-RS* signals were predominantly derived from osteoblastic OS cells and CAFs (Fig. 3B).

**Fig. 3:**
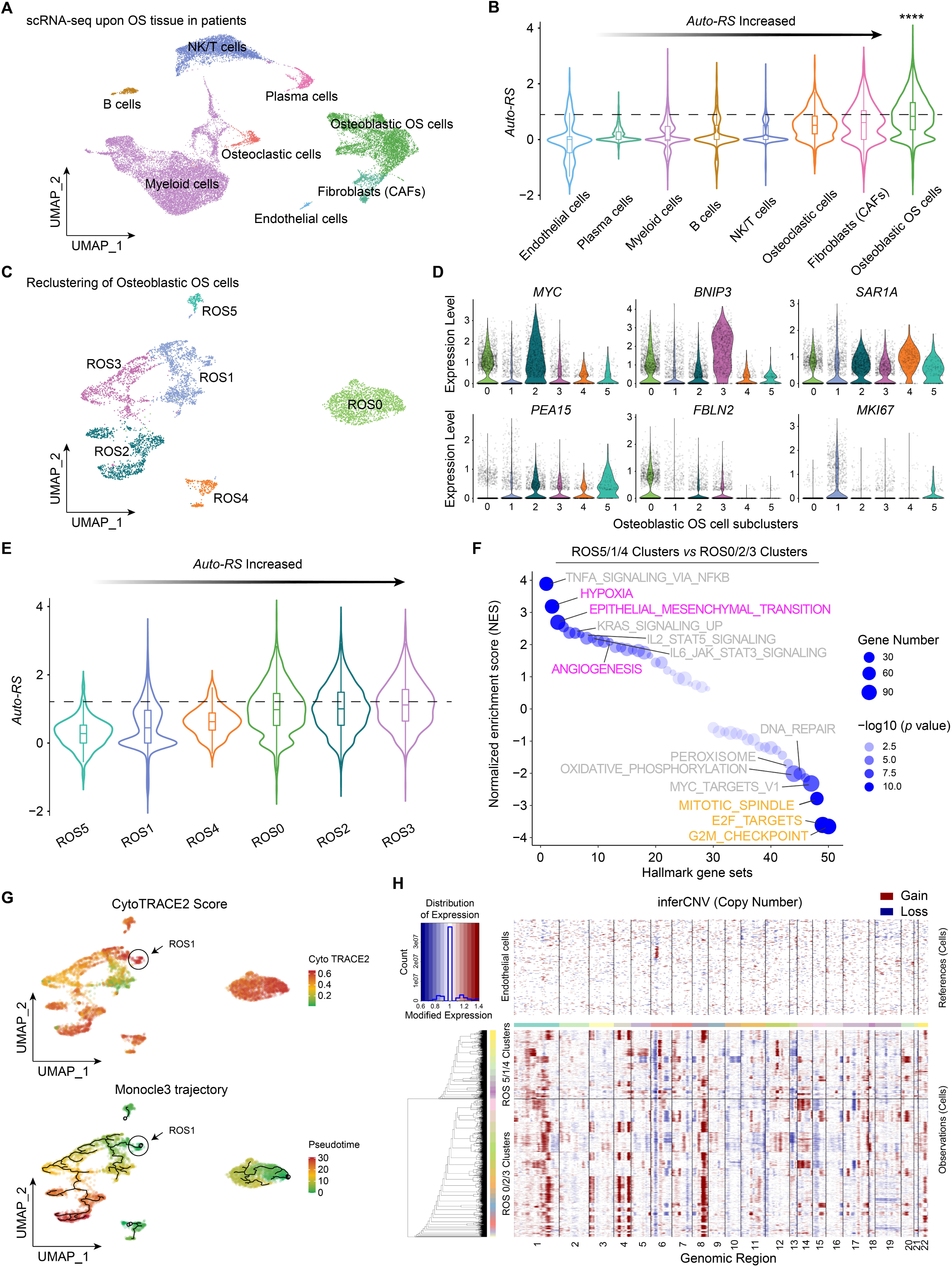
Single-cell analysis reveals the cellular origin and molecular basis of *Auto-RS* prognostic signals. **A**, Uniform manifold approximation and projection (UMAP) plot embedding of 27,719 cells from six primary OS tissues delineating eight major populations. **B**, *Auto-RS* distribution across the eight major cell populations, increasing gradually from left to right. **C**, Reclustering of 5,988 osteoblastic OS cells identified six subclusters (ROS0 to ROS5). **D**, Violin plots showing the expression of *Auto-RS* related genes, including differential expression of *FBLN2* and *MKI67*. **E**, The *Auto-RS* signal was mainly enriched in ROS0/2/3. **F**, GSEA of Hallmark pathways contrasting ROS0/2/3 (purple) with ROS5/1/4 (yellow), revealing enrichment of hypoxia, EMT, angiogenesis, and KRAS signaling versus proliferative signatures (G2M checkpoint, E2F targets, mitotic spindle). Circle size reflects gene counts and color intensity indicates significance. **G**, CytoTRACE2 and Monocle3 analysis depicting differentiation potential and pseudotime trajectories. High-stemness, undifferentiated ROS1 cells evolve along distinct branches toward metastasis-associated *Auto-RS^high^* ROS2/3. **H**, Clustering of copy number variation (CNV) profiles inferred from scRNA-seq data for osteoblastic OS cells and endothelial cells. Clusters (dendrogram) primarily reflect ROS0/2/3 and ROS5/1/4 CNVs (colored bar coded). The CNV signal heatmap, normalized against the ‘normal’ cluster defined by endothelial cells, shows chromosome-wise CNV alterations (columns) across individual cells (rows).

Within 5,988 osteoblastic cells, *Louvain* clustering identified six subpopulations (ROS0–5) with heterogeneous *Auto-RS* expression. (Fig. 3C and S3D). *Auto-RS* was elevated in ROS0/2/3 relative to ROS5/1/4, reflecting differential expression of the four autophagy genes (Fig. 3D and 3E). Functionally, ROS0/2/3 were characterized by metastatic programs, with enrichment of adhesion molecules such as *FBLN2*, along with hypoxia, EMT, angiogenesis, and KRAS signaling^23^ (Fig. 3E and 3F, ***Table S9***). KEGG and GO analyses further highlighted activation of skeletal development (ossification, bone and cartilage formation), stress-adaptive pathways (hypoxia, unfolded protein response), and canonical oncogenic cascades including TGF-β, MAPK, PI3K-AKT, FoxO, Wnt, HIF-1, and Relaxin. *Auto-RS*^high^ clusters also upregulated stemness signatures, extracellular matrix (ECM)-receptor interactions, and focal adhesion, consistent with enhanced self-renewal and invasive plasticity (Fig. S3E-H). By contrast, *Auto-RS*^low^ ROS5/1/4 displayed proliferative states, marked by *MKI67* expression, enrichment of cell cycle and mitotic pathways, and metabolic activation including oxidative phosphorylation and ATP biosynthesis (Fig. 3E, F, and S4A-D).

Trajectory inference using CytoTRACE2^24^ and Monocle3^25^ revealed less-differentiated, highly proliferative ROS1 cells occupy early pseudotime states with low *Auto-RS*, whereas progression toward high *Auto-RS* ROS2/3 coincides with acquisition of metastasis-associated transcriptional programs (Fig. 3G). Early transitions involved stress response and proteostasis genes (*HSPA5*, *HSP90AA1*, *BAG3*, *ATF3*), intermediate stages showed elevated ribosomal protein expression (*RPL30*, *RPL32*, *RPS3A*), and terminal stages activated inflammatory, anti-apoptotic, and cytoskeletal remodeling modules (*FOS*, *JUN*, *MCL1*, *FN1*), defining a coherent high *Auto-RS* metastatic program (Fig. S4E).

Finally, inferred CNV^26^ analysis revealed that *Auto-RS* differences were largely independent of global chromosomal alterations (Fig. 3H). ROS0/2/3 and ROS5/1/4 subpopulations exhibited similar overall CNV burdens (Fig. S5A and S5B), with only *CDK4* amplification in ROS5/1/4 may partially explaining their proliferative phenotype. Other recurrent OS CNVs and targetable signaling genes (*MYC*, *PTEN*, *WWOX*, *FLI1*, *NOTCH2*, *PDGFRA*, *VEGF*, *IGF1R*)^27,28^ did not correlate with *Auto-RS* stratification (Fig. S5C).

Together, these data indicate that *Auto-RS* signals originate primarily from osteoblastic OS cells, with *Auto-RS*^high^ subpopulations exhibiting stress- and metastasis-related programs, while *Auto-RS*^low^ cells are highly proliferative, bioenergetically active states. *Auto-RS* thus captures dynamic cellular states linked to lineage plasticity and metastatic potential beyond genomic alterations.

### *Auto-RS* permeate the metastatic OS microenvironment

Efficient metastatic progression is tightly orchestrated through dynamic cross-talk among tumor cells, cancer-associated fibroblasts (CAFs), and immune populations^29^. CAFs exhibited strong *Auto-RS* activity and segregated into four functional subtypes: matrix CAFs (mCAFs), vascular CAFs (vCAFs), antigen-presenting CAFs (apCAFs), and inflammatory CAFs (iCAFs)^30^ (Fig. 4A and 4B). Although vCAFs and apCAFs were numerically dominant (Fig. 4C), mCAFs contributed the highest *Auto-RS* signal (Fig. 4D), characterized by ECM-remodeling and pro-metastatic mediators (*IL11*^31^, *TGFB1*, *INHBA*, *MIF*, *PTHLH*). In contrast, vCAFs were enriched for angiogenesis drivers (*NOTCH1–3*, *JAG1*, *ANGPT2*^32^, *PDGFA*, *MMP9/11*^33,34^), iCAFs secreted inflammatory and chemoresistance-associated cytokines (*IL6*, *LIF*, *IGF1*, *CCL2/11*), with *HAS1* upregulation linked to hyaluronan-mediated metastasis^35,36^, and apCAFs combined antigen presentation with immune-exclusion programs (*CXCL12*, *THBS1*, *MDK*) (Fig. 4E and 4F, ***Table S10***).

**Fig. 4:**
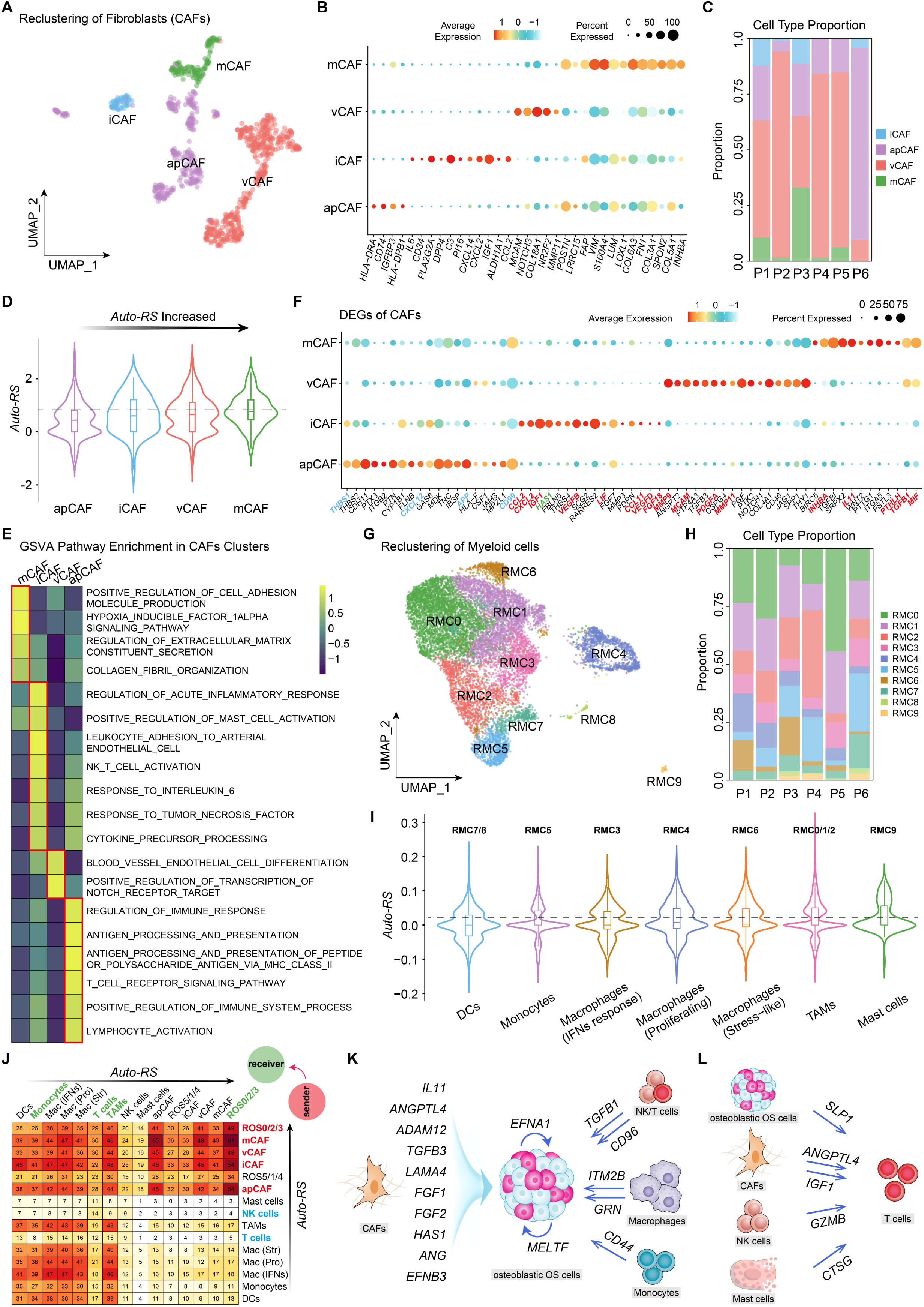
Single-cell dissection of *Auto-RS* signals across CAF and immune populations. **A**, Reclustering of CAFs revealed four distinct subsets: matrix CAFs (mCAFs), vascular CAFs (vCAFs), inflammatory CAFs (iCAFs), and antigen-presenting CAFs (apCAFs). **B**, Dot plot of representative marker genes across CAF subsets. Dot size indicates the proportion of expressing cells; color intensity reflects relative expression levels. **C**, Stacked bar chart showing the distribution of CAF subsets across six patients, with vCAFs (48.6%) and apCAFs (31.3%) constituting the majority, followed by iCAFs (14%) and mCAFs (6.1%). **D**, Distribution of *Auto-RS* values across CAF subsets, ranked from low to high. **E**, Heatmap of normalized GSVA scores for selected Gene Ontology pathways in each CAF subset. **F**, Dot plot showing DEGs across mCAF, vCAF, iCAF and apCAF clusters. **G**, Reclustering of myeloid cells identified 10 subtypes (RMC0 to RMC9). **H**, Stacked bar chart showing the proportion of RMC0 to RMC9 clusters in six OS patients. **I**, Distribution of *Auto-RS* values across myeloid subsets, ranked from low to high. **J**, Number of ligand–receptor interactions among osteoblastic OS subtypes, CAF subsets, myeloid subsets, NK cells, and T cells. Rows represent sender cells (ordered from low to high *Auto-RS*, bottom to top); columns represent receiver cells (ordered from low to high *Auto-RS*, left to right). **K**,**L**, *Auto-RS* identified key pro-metastatic cell — cell communications within the OS microenvironment, including signals promoting osteoblastic OS cell metastasis (**K**) and suppressing T-cell antitumor immunity (**L**).

Before analyzing cross-talk, myeloid cells were further resolved into mast cells, dendritic cells (DCs), monocytes, tumor-associated macrophages (TAMs), and three specialized macrophage states (proliferating, stress-like, IFN-responsive) (Fig. 4G-I). Notably, neutrophil markers (S100A8/9) localized within the monocyte cluster, precluding neutrophils as a distinct population (Fig. S6A and S6B). *CellChat* program^37^ revealed that osteoblastic OS cells and CAFs with high *Auto-RS* were dominant “signal senders,” engaging in dense reciprocal interactions with each other and with immune cells. By contrast, immune populations with low *Auto-RS* contributed relatively few outgoing signals (Fig. 4J and S7A, ***Table S11***). *NicheNet* program^38^ highlighted CAF-derived pro-metastatic ligands as key regulators of *Auto-RS*^high^ tumor cells, including *ANGPTL4/ADAM12* (mCAFs), *TGFB3/LAMA4/FGF1* (vCAFs), *FGF2/HAS1/ANG* (iCAFs), and *EFNB3*^39^ (apCAFs) (Fig. S7B). ROS0/2/3 cells reinforced this network via autocrine *BMP2/SEMA3A/EFNA1*, with *MELTF* from ROS5/1/4 further augmenting its metastatic potential. Unexpectedly, NK cells—rather than mCAFs—emerged as a major source of *TGFB1*, a canonical EMT driver (Fig. S7C). In parallel, NK/T cells transmitted inhibitory signals through *CD96*^40^, suppressing their own cytokine responses and cytotoxic activity. TAMs and monocytes further promoted metastasis through *ITM2B/GRN* and *CD44/S100A4/VSTM1/NAMPT*, respectively (Fig. 4K).

Although T cells contributed minimally to the *Auto-RS* signal itself, their functional states were sensitively captured. Stratification revealed that *Auto-RS*^high^ T cells received *SLP1*^41^ from ROS0/2/3, a serine protease inhibitor that impairs cytotoxic T lymphocyte function by blocking lytic granule release, thereby enhancing tumorigenic and metastatic potential. In addition, mCAFs and iCAFs suppressed T-cell immunity via *ANGPTL4*^42^ and *IGF1*^43^, respectively. *Auto-RS*^high^ T cells also received proteolytic inhibitory inputs from mast cells (*CTSG*) and NK cells (*GZMB*), both of which can severely compromise T-cell activity (Fig. 4L).

Together, these findings delineate a pervasive *Auto-RS* driven signaling network in which CAF subsets and malignant OS cells orchestrate immune suppression and metastatic competence, while *Auto-RS*^high^ immune cells acquire impaired antitumor functionality.

### *Auto-RS* stratification identifies therapeutic vulnerabilities

To explore whether *Auto-RS* stratification uncovers druggable dependencies^44^, we applied it to *Precily*^45^, a deep learning–based pharmacogenomic predictor (Fig. 5A). Predicted responses of 139 OS patients to 156 compounds tested across four OS cell lines (HOS, MG63, U2OS, SAOS2) (***Table S12***) identified nine agents with the highest sensitivity: microtubule-targeting drugs (vinorelbine, vincristine, vinblastine), apoptosis regulators (staurosporine, sepantronium bromide, obatoclax), DNA synthesis inhibitors (gemcitabine, cytarabine), and the FGFR inhibitor AZD4547. In contrast, EGFR/ERBB inhibitors were consistently ineffective, in line with prior reports^46^.

**Fig. 5:**
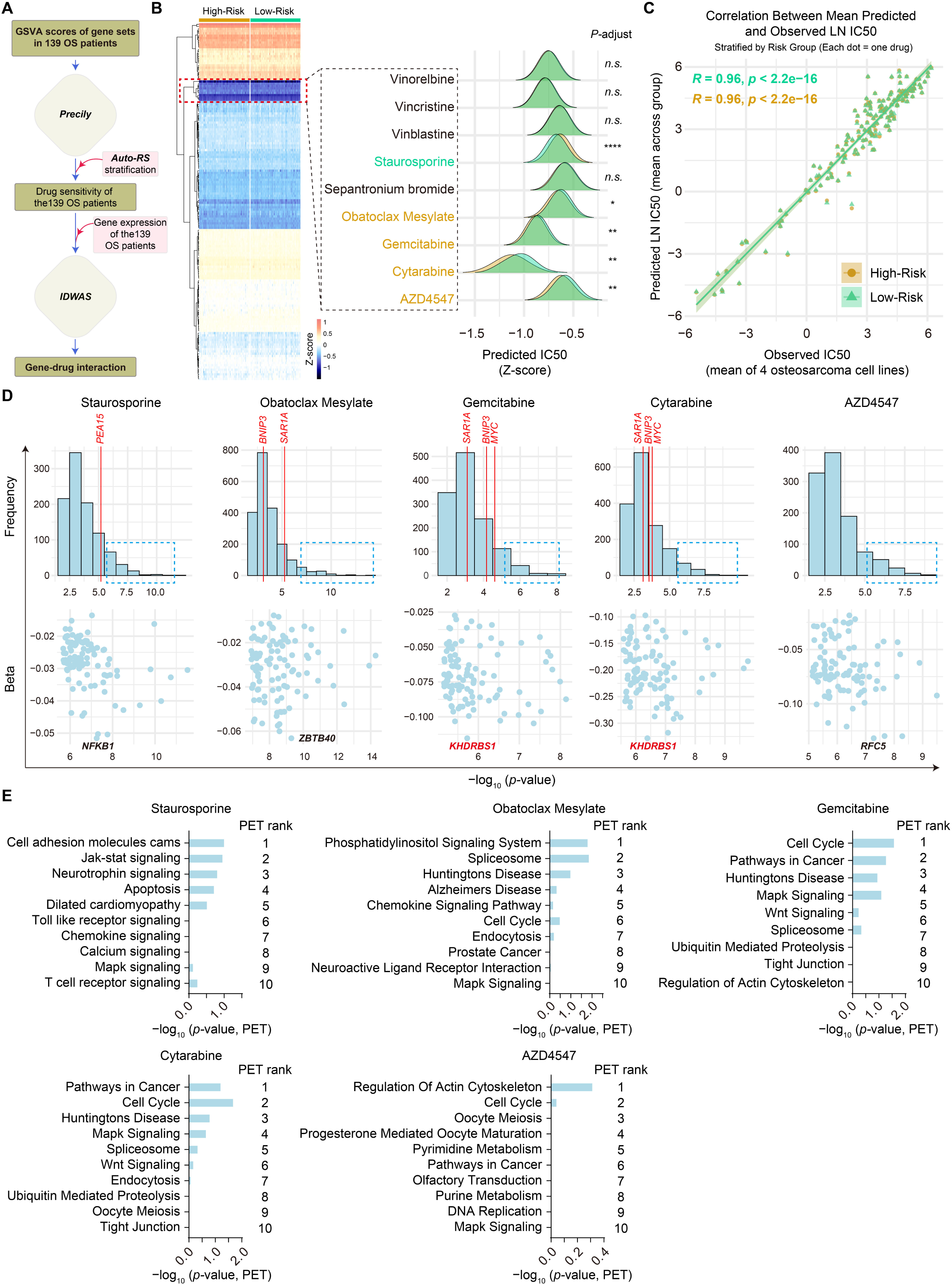
Therapeutic vulnerabilities identified through *Auto-RS* based drug sensitivity prediction. **A**, Workflow for predicting drug sensitivity and identifying potential therapeutic targets using *Auto-RS* stratification. **B**, Heatmap of predicted IC50 Z-scores for 156 compounds across four OS cell lines (HOS, MG63, U2OS, SAOS2) in high- and low-risk groups. Lower IC50 indicates higher predicted sensitivity. Ridge plots show the distribution of predicted IC50 for the nine most sensitive drugs, with green indicating higher sensitivity in low-risk and yellow in high-risk groups. *P*-values were calculated by Wilcoxon rank-sum test (< 0.05*, < 0.01**, < 0.0001***, *n.s.*, not significant). **C**, Scatter plot showing the correlation between predicted and observed IC50 values (Pearson’s *R*) across high- and low-risk groups. Two-sided t-test was used for *P*-value calculation; shaded areas indicate 95% confidence intervals. **D**, IDWAS analysis linking patient gene expression profiles to predicted drug response. Top panel shows the frequency of significant gene–drug associations (*P*-values) across all genes. The blue dashed box highlights the top 100 genes with the smallest *P*-values, displayed in the scatter plot below. Beta coefficients indicate the effect of gene expression on predicted IC50, with more negative values denoting higher sensitivity. *P*-values were computed using two-sided t-tests. **E**, Pathway enrichment analysis of target genes corresponding to five predicted sensitive drugs using the Pathway Ensemble Tool (PET). *P*-values represent combined significance estimated via the Stouffer method integrating GSEA, ORA, and Enrichr results. Top 10 pathways are ranked by average score across methods.

Median Z-scores derived from predicted IC50 values identified five compounds with *Auto-RS* specific vulnerabilities: staurosporine showed preferential efficacy in the low-risk group, whereas obatoclax, gemcitabine, cytarabine, and AZD4547 were more effective in high-risk group (Fig. 5B). Notably, gemcitabine and cytarabine are FDA-approved agents for solid tumors and leukemia, respectively. Gemcitabine has demonstrated tolerability in OS combination regimens^47,48^, whereas cytarabine has not yet been clinically evaluated in OS, despite preclinical sensitivity in methotrexate-resistant cell lines^49^. Predicted IC50 values correlated strongly with observed IC50s across GDSC cell lines (Pearson’s *R* = 0.96, *P* < 2.2 × 10^-16^; Fig. 5C), validating the robustness of the predictions. Refined imputed drug–wide association (IDWAS)^50^ further identified predictive biomarkers, as the number of clinically validated biomarkers for osteosarcoma has been described as “strikingly small”^11^ (Fig. 5D, and ***Table S13***). Sensitivity to staurosporine in low-risk patients was linked to *NFKB1* expression, whereas *ZBTB40* predicted response to obatoclax. *RFC5* emerged as a marker of AZD4547 sensitivity. For the FDA-approved agents, gemcitabine and cytarabine responses converged on *KHDRBS1*^51^ as a shared, highly specific biomarker, consistent with its established role in chemosensitivity of cancer stem cells. Pathway Ensemble Tool (PET)^52^ highlighted *cell cycle* regulation as the principal pathway perturbed by gemcitabine and cytarabine (Fig. 5E).

Together, these results establish *Auto-RS* stratification as a framework for uncovering therapeutic vulnerabilities and predictive biomarkers in OS, with particular relevance for high-risk patients.

## Discussion

In this study, we asked whether autophagy programs could be leveraged as precise readouts of metastatic biology, offering a paradigm for clinically actionable prognostic modeling in aggressive tumors. The LASSO-Cox regression–based machine learning framework was applied to datasets from 139 patients, integrating autophagy-related transcriptional features with clinical variables to derive the *Auto-RS* signal and construct the ApM. The ApM achieved a C-index of 0.709 for early risk stratification and further enhanced individualized prognostic management when integrated with metastasis information. We ensured that models were trained on the currently most comprehensive clinical and transcriptomic OS datasets and then evaluated across independent single-cell datasets. Confronted with expression landscapes encompassing 13,770-gene profiles, the *Auto-RS* classifier distilled metastasis-specific patterns and prioritized a minimal yet informative gene set, which in combination with age and sex, yielded robust prognostic predictions. *Auto-RS* generalized to independent single-cell sequencing data from other laboratories, served as a conserved molecular barcode capable of penetrating the complex tumor microenvironment, capturing metastatic competence across tumor cell and stromal compartments. This deep conservation of *Auto-RS* suggests that its upstream regulation may represent a pivotal checkpoint in the malignant evolution of tumors.

Key innovations underlying *Auto-RS* performance stem from iterative self-supervised screening of autophagy programs within the validation pipeline, combined with independent single-cell functional interpretation. These two components, though distinct, are complementary; the integrated validation approach outperformed single-aspect analysis. A representative case was that models constructed on bulk transcriptomes or conventional metastasis-associated transcripts exhibited limited interpretability in single-cell analyses owing to noisy signal origins (Fig. S8). For example, *Auto-RS* appeared to derive predominantly from B cells and plasma cells. Likewise, *Auto-RS* displayed restricted discriminatory power across cellular subpopulations, a result at odds with the marked heterogeneity of the OS microenvironment^53^. Thus, independent interpretability validation proved essential; incorporating both the cellular origins of signals within the tumor microenvironment and single-cell functional information yielded a more comprehensive understanding of prognostic states.

*Auto-RS* delineated the cell-intrinsic functional changes that contributed most to prognostic predictions in OS. We confirmed that osteoblastic OS cells and CAFs, previously reported in the literature as key drivers of metastasis^53^, carried the greatest weight in the *Auto-RS* prediction process. This would be consistent with *Auto-RS* learning biologically meaningful transcriptional features rather than simply fitting to dataset-specific expression changes. Unlike studies restricted to a specific cell type^54,55^, *Auto-RS* enables pinpointing metastasis-promoting responses unique to each cellular compartment. Prior to extensive experimental validation, *Auto-RS* may thus serve as a guiding framework to simultaneously distinguish critical pro-metastatic cell types and their signaling molecules, facilitating the discovery of therapeutic targets and candidate drugs. Our standardized bioinformatic analyses combined with *Auto-RS* attempted to address these challenges, highlighting several molecular targets (*IL11, HAS1, ANGPT4, SLP1, IGF1*) and the drug cytarabine as potential vulnerabilities.

*Auto-RS* is biologically grounded, reflecting autophagy-mediated transcriptional “stress fingerprints” that are tightly linked to metastasis. Functionally, *BNIP3* promotes anoikis resistance during ECM detachment^56,57^, *SAR1A*-mediated secretory autophagy links ER stress to metastasis^58,59^, *MYC* drives ER stress–induced autophagy to facilitate malignant transformation^60^, whereas *PEA15* exerts tumor-suppressive effects^61,62^. These mechanistic insights indicate that *Auto-RS* captures conserved cellular stress responses within tumor cells rather than random transcriptional noise. Although the roles of these genes in stromal and immune compartments remain less well defined, *Auto-RS* may be best interpreted as a composite readout of stress-adaptive states across the tumor ecosystem, reflecting the pre-metastatic condition rather than representing a direct causal program driven solely by transcriptional dysregulation of these four individual genes.

Our work underscores the value of deep transcriptomic profiling for unlocking prognostic signals, even from limited patient material. Because transcriptional changes often precede phenotypic manifestations, *Auto-RS* can capture metastatic potential more sensitively than conventional physiological readouts. Nevertheless, several limitations remain. It is unclear whether *BNIP3, SAR1A, MYC,* and *PEA15* directly drive metastasis or instead mark pre-metastatic states; the latter is more likely, as it is improbable that four autophagy-related genes alone could orchestrate consistent functional transitions across diverse cell types. Furthermore, additional prospective studies will be needed to define clinically meaningful thresholds for sensitivity and specificity, optimize sample size, and evaluate sequencing depth. Results from this approach should be interpreted alongside other clinical assessments and laboratory testing.

## Methods

### Patient data acquisition and quality control

RNA sequencing data matrices for TARGET were obtained from the Genomic Data Commons (https://portal.gdc.cancer.gov/). These matrices integrate both gene expression profiles and clinical characteristics of osteosarcoma patients. Genes with zero expression were removed, and duplicate genes were merged by averaging their expression values. The processed data were subsequently log2-transformed. In addition, the expression matrix and corresponding clinical information of the GSE21257 cohort were downloaded from the Gene Expression Omnibus (GEO, https://www.ncbi.nlm.nih.gov/geo/). Preprocessing followed the same pipeline, including the removal of genes with zero expression and the averaging of duplicate genes. Patients lacking complete clinical information (follow-up time, age, sex, metastasis status, or survival outcome) were excluded. After quality control, the TARGET cohort comprised 86 OS samples with 56,515 genes, whereas the GSE21257 cohort included 53 OS samples with 24,970 genes.

### Autophagy gene extraction

A total of 222 autophagy-related genes were obtained from the Human Autophagy Database (HADb, https://www.autophagy.lu/v1/) (***Table S2***). Of these, 222 autophagy genes were extracted from the TARGET cohort and 193 from the GSE21257 cohort, with 193 autophagy genes shared across both datasets. The intersection of autophagy genes was visualized using a Venn diagram generated with the R package *VennDiagram* (v1.7.3).

### Batch effect correction and identification of differentially expressed genes

To correct for batch effects, expression matrices from TARGET and GSE21257 were combined and processed using the *sva* package (v3.54.0) together with the limma::removeBatchEffect function. In addition, *BatchServer* (https://lifeinfo.shinyapps.io/batchserver/), an online tool integrating principal component analysis (PCA) implemented in the *factoextra* package (v1.0.7), was applied to eliminate and assess batch effects. The results were visualized with pie charts and PCA plots. The merged cohort consisted of 193 autophagy genes (***Table S3***) across 139 OS samples and was used for subsequent analyses. Differential expression analysis between metastatic and non-metastatic groups was assessed using the *limma* package (v3.62.2), with the thresholds set as |log2FoldChange| > 0.2 and *P* < 0.05. Heatmaps and volcano plots were generated to visualize DEGs using the R packages *pheatmap* (v1.0.12), *RColorBrewer* (v1.1.3), and *ggplot2* (v3.5.1).

### Construction and validation of the prognostic risk model

To construct a robust prognostic model, we integrated the expression matrix of DEGs with patient survival data and employed an iterative feature selection strategy combined with a dual-validation mechanism to ensure model stability. To balance feature selection stringency with interpretability, we adopted a hybrid approach that integrates the Least Absolute Shrinkage and Selection Operator (LASSO) with bidirectional stepwise regression, which optimizes the model by minimizing the AIC.

The iterative procedure was designed as follows:

I. Data Partitioning: (1) The dataset was randomly split into training and testing sets in a 1:1 ratio using the createDataPartition function. (2) This randomization was repeated 1,000 times to ensure model stability.
II. Feature Selection: (1) Univariate Cox regression was performed to assess the association between gene expression and overall survival (*P* < 0.05), retaining only those genes that were statistically significant in the training set. (2) LASSO regression: (i) Performed with 10-fold cross-validation to determine the optimal λ value. (ii) Genes with non-zero coefficients at lambda.min were retained. (iii) If no gene was selected, the iteration was skipped.
III. Multivariate Cox Regression Model Construction: (1) The selected features were subjected to bidirectional stepwise regression. (2) The *Auto-RS* was calculated using the following formulation:

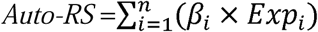

Where β*_i_*is the Cox regression coefficient and Exp*_i_* is the expression level of gene *i*.
IV. Model Validation: (1) Validation was conducted in both the training and testing sets: (i) Survival difference was evaluated using the log-rank test (*P* < 0.05). (ii) ROC curve analyses were performed, and models with AUC > 0.75 were considered acceptable. (2) Iterations were terminated when both datasets met the evaluation criteria simultaneously. Following the above pipeline, 20 candidate models were retained (***Table S5***). For each model, samples in both the TARGET and GSE21257 cohorts were divided into high-risk and low-risk groups based on the median *Auto-RS* for further validation. Groups with statistically significant survival differences (*P* < 0.05) in both cohorts were retained, resulting in 8 validated models (***Table S6***).

To optimize model complexity and performance, we prioritized models with the best balance of fit and simplicity using stepwise regression and assessed their discrimination ability using the concordance index (C-index). The steps for model selection are as follows:

I. Compute the ΔAIC (AIC value minus the minimum AIC across models).
II. Exclude models with ΔAIC > 10.
III. Among the remaining models, rank them by C-index in descending order and select the one with the highest C-index.

Through this process, the best iteration was ultimately selected as the optimal prognostic model for downstream analyses (***Table S7***).

For model construction, we used the *survival* package (v 3.8-3) to perform univariate Cox regression and identify autophagy gene significantly associated with overall survival (*P* < 0.05). We employed the *glmnet* package (v 4.1.8) to run 1,000 LASSO iterations, obtaining robust regression coefficients with 10-fold cross-validation. The final multivariate Cox model was built using the “coxph” function combined with stepwise regression to derive the optimal prognostic signature.

Model performance was visualized using Kaplan–Meier survival curves and time-dependent ROC curves were generated using *survival* (v3.8.3), *survminer* (v0.5.0), and *timeROC* (v0.4), enabling assessment of sensitivity and specificity. A risk heatmap for the four model genes was generated using *pheatmap* (v 1.0.12). Patient *Auto-RS* distribution plots and survival status scatter plots were generated using *ggplot2* (v 3.5.1).

To evaluate whether the *Auto-RS* functioned as an independent prognostic factor, univariate and multivariate Cox regression analyses were performed using the *survival* package (v3.8-3).

Finally, nomograms were constructed to visualize the prognostic model components. Three types of models were defined:

I. Autophagy prediction model (ApM): including *Auto-RS*, Age, and Gender.
II. Basic prediction model (BpM): including Metastases, Age, and Gender.
III. Complete prediction model (CpM): including *Auto-RS*, Metastases, Age, and Gender.

In the nomograms, the age variable was modeled using a restricted cubic spline function, with three knots (rcs, k = 3) to capture potential non-linear associations with the outcome variable. Quantified scoring tables were generated ***TableS1***. The *rms* package (v 7.0.0) was used to compute calibration curves for evaluating prediction reliability—models with calibration curves closer to the diagonal line indicate better predictive accuracy. Additionally, the C-index was calculated to evaluate the predictive ability of each nomogram.

### Single-cell RNA sequencing data of OS patients

Raw scRNA-seq data of six primary OS tissues were obtained from the GEO database (GSE162454^22^). Downstream analyses were run with *Seurat* (v5.2.1) package in R (v4.4.2). For each single-cell RNA-seq data, we created a Seurat object with the CreateSeuratObject function. All *Seurat* objects were merged to one and filtering steps as follows were applied for the merged data: (1) cells with fewer than 200 detected genes, more than 6000 detected genes, or with UMI counts exceeding 30000 were excluded; (2) cells with percentage of mitochondrial UMI count larger than 10% were filtered out; (3) the genes detected in more than 3 cells were retained and all ribosomal genes were filtered out. After quality control, a total of 27,719 high-quality cells were retained. Since the samples of OS tissues come from different patient individuals, this may introduce technical and biological batch effects in downstream analysis. Therefore, based on PCA, the RunHarmony function of *harmony* (v1.2.3) R package was used to correct the batch effect.

### Cell type identification and clustering

Initial sample-level clustering was performed in three sequential steps: (1) dimensionality reduction by principal component analysis (PCA) using all genes as variable features; (2) construction of a shared nearest-neighbor graph in PCA space using the FindNeighbors function; (3) unsupervised clustering using the FindClusters function (unsupervised clustering was performed using the shared nearest neighbor (SNN) modularity optimization algorithm with the Louvain method) at a resolution of 0.2. DEGs analysis was performed using the FindAllMarkers function in the *Seurat* package. Significant DEGs were identified using a cutoff of an FDR-adjusted *P*-value < 0.05 and a log2 fold change > 0.25. Cell type annotation was performed based on canonical marker gene expression, as follows: Myeloid cells: *LYZ, CD68, CD163*.Cancer-associated fibroblasts (CAFs): *FBLN1, ACTA2, TAGLN, COL3A1, COL6A1*. Osteoblastic OS cells: *ALPL, RUNX2, IBSP*. Osteoclastic cells: *ACP5, CTSK*. B cells: *CD79A, MS4A1*. NK/T cells: *CD2, CD3D, CD3E, CD3G, GNLY, NKG7, KLRD1, KLRB1*. Endothelial cells: *EGFL7, PLVAP.* Plasma cells: *IGHG1, MZB1*. For sub-clustering, the subset function was used to extract specific cell populations. Reclustering was then performed following the same pipeline for osteoblastic OS cells (resolution = 0.1, *n* = 5,988), CAFs (resolution = 0.1, *n* = 915), and myeloid cells (resolution = 0.3, *n* = 12,876).

### Pathway Analysis

Gene Ontology (GO) and Kyoto Encyclopedia of Genes and Genomes (KEGG) enrichment analyses of DEGs were performed using the *clusterProfiler* package (v4.14.6). Gene Set Variation Analysis (GSVA) was conducted with the *GSVA* package (v2.0.7).

### Copy Number Variation Analysis

To assess copy number variations (CNVs) in tumor cells across clusters, we performed CNV analysis using the *inferCNV* package (v1.23.0). Endothelial cells were designated as the reference population, defined as reference Endothelial, to provide a baseline for CNV inference. Comparative analyses were conducted against clusters ROS0/2/3 and ROS5/1/4 to calculate the variability of gene expression intensity across chromosomes in Osteoblastic OS cells. The key parameters were set as denoise = TRUE, HMM = FALSE, and cutoff = 0.1, with all other parameters left at their default values. Furthermore, to facilitate interactive exploration of specific genes and chromosomal regions, we utilized the next-generation clustered heatmap (NG-CHM) tool^27^. The processed infercnv.ngchm data can be interactively visualized and explored at https://www.ngchm.net/Downloads/ngChmApp.html.

### Prediction of differentiation states

We used *CytoTRACE2*^24^ package (v1.1.0) to predict the differentiation states of osteoblastic OS cells in our scRNA-seq data. This method provides a continuous assessment of developmental potential through predicted potency scores ranging from 0 to 1, where 0 represents differentiated cells and 1 corresponds to totipotent cells. The raw count matrix was input to generate the prediction using the CytoTRACE2 function, and dedifferentiation scores were plotted with the plotData function using UMAP coordinates generated from *Seurat*.

### Pseudotime trajectory analysis

To explore the developmental trajectory of OS cells, we used *Monocle3*^25^ (v1.4.26) to order cells along a pseudotemporal axis based on transcriptional similarity. A trajectory graph was constructed using the learn_graph function, with the top 1% of cells exhibiting the highest stemness scores in CytoTRACE2 selected as the starting point for pseudotime analysis. To visualize cell distribution and gene expression dynamics along pseudotime, heatmaps of gene expression were generated using the *pheatmap* package (v1.0.13).

### CellChat analysis

We used CellChat^37^ (v1.6.1) for the analysis of cell–cell communication and the analysis followed the official workflow with default parameters unless otherwise specified. First, we loaded the normalized counts into CellChat, followed by the preprocessing steps identifyOverExpressedGenes and identifyOverExpressedInteractions. We then ran the computeCommunProb function for communication analysis with the parameter population.size = FALSE to eliminate possible bias due to cell population size. This resulted in a network of communication strength between all cell states for each of the ligand–receptor pairs that passed the filtering steps. We used the aggregation functions computeCommunProbPathway and aggregateNet to determine the communication strength between cell states at pathway and global levels, respectively. In addition, we applied the *NicheNet*^38^ method to perform a detailed analysis of predicted ligands driving expression changes in target cell subpopulations. Target cell subpopulations of interest included ROS0/2/3 and T cell subsets, while all cells from the relevant sender subpopulations were considered as potential ligand sources. Default parameters were used to predict intercellular interactions.

### Drug Response Prediction

We employed the *Precily* algorithm^45^, a machine learning framework for predicting drug responses based on gene expression data. In this approach, pathway enrichment scores derived from RNA-seq expression data via GSVA are integrated with molecular descriptors of drugs (converted from SMILES to vector representations) to form a unified feature matrix. Regression modeling is performed using multiple algorithms, including Random Forests (implemented with the *ranger* package), ElasticNet (implemented with the *glmnet* package), and deep neural networks (implemented with the *Keras* framework). In the original study, the authors trained and validated the model using CCLE and TCGA datasets, optimized hyperparameters via *k*-fold cross-validation, and ultimately predicted drug sensitivity (IC50 or response classification).

In this study, the data processing pipeline recommended by the *Precily* was followed, and the required input files were prepared as follows:

I. Pathway enrichment scores: Based on log (TPM + 1) gene expression matrices from osteosarcoma samples and cell lines, Gene Set Variation Analysis (GSVA) scores were calculated using the *GSVA* package (v2.0.7) in R, with min.sz = 5. Gene sets were sourced from the c2 canonical pathway collection in the MSigDB database (MSigDB.CP.v6.1), comprising 1,329 gene sets, consistent with the original model training set.
II. Drug and sample metadata: Metadata files were initially provided by the authors, and we supplemented them with annotations for osteosarcoma cell lines (HOS, MG63, U2OS, and SAOS2) based on the Genomics of Drug Sensitivity in Cancer (GDSC) database. The two input files were then passed to the Precily drugPred function as follows: drugPred (enrichment.scores, metadata, "OS"). This process generated predicted drug sensitivities (IC50 values) for 139 osteosarcoma patient samples and 4 osteosarcoma cell lines across 156 drugs.

### Prediction of Potential Drug Biomarkers

To discover potential biomarkers of drug sensitivity, we utilized the imputed drug-wide association study (IDWAS) method^50^. This approach employs linear ridge regression to identify associations between predicted drug sensitivity and gene expression data. We integrated the predicted drug response data with the gene expression matrix of 139 osteosarcoma samples to systematically evaluate gene–drug correlations.

### Pathway Prediction for Potential Drug Biomarkers

To further investigate the biological pathways potentially involved in drug biomarker genes, we employed the Pathway Ensemble Tool (PET)^52^. PET integrates three widely used pathway enrichment approaches (GSEA, ORA, and Enrichr) to calculate independent enrichment scores for each pathway and subsequently generates a consensus ranking across methods. We then derived the average rank across the three approaches and applied the Stouffer method to estimate a combined *P* - value. Finally, pathways were ranked according to their aggregated scores, enabling the identification of biological mechanisms potentially associated with drug sensitivity.

### Statistics and reproducibility

No statistical method was used to predetermine sample size. All statistical analyses were performed in *R* (v4.4.1). Survival curves were generated using the Kaplan–Meier method and compared with the log-rank test. Comparisons between two groups were conducted using either the unpaired two-tailed Student’s t-test or the Wilcoxon rank-sum test, as appropriate. The Likelihood Ratio Test was employed to assess the statistical significance of differences in c-index between models. The Wald test was used to test the significance of univariate and multivariate Cox models. Pearson and Spearman correlation coefficients were used to evaluate associations between continuous variables. *P* values less than 0.05 were considered statistically significant.

### Code availability

All code necessary for the analyses is available without access restrictions via GitHub at: https://github.com/Lqz-DC/Autophagy-in-osteosarcoma.git. All data generated or analyzed during this study are available from the corresponding author upon reasonable request.

## Acknowledgements

This work was supported by grants from National Natural Science Foundation of China (grant nos. 82201533), National Natural Science Foundation Incubation Program of The Second Affiliated Hospital of Anhui Medical University (2021GQFY02), and Anhui Medical University Foundation (9101224201). The funders had no role in the study design, data collection and analysis, decision to publish or preparation of the paper.

## Author contributions

Conceptualization: Y.B., Q.L. and J.L.; data curation: Q.L., C.Z., K.X., and J.L.; formal analysis: Y.B., Q.L., C.Z., K.X., Q.Z., and J.L.; funding acquisition: Y.B.; methodology: Y.B., Q.L., Q.Z., and J.L.; project administration: Y.B., W.D., and H.W.; software: Q.L., C.Z., K.X., Q.Z., and J.L.; validation: C.Z., G.F., X.C.; data collection: Q.L., C.Z., K.X., and J.L.; data quality control: Q.Z., W.X., D.T., and J.J.; writing—original draft: Y.B., Q.L., W.D., and H.W.; writing—review and editing: Q.Z., Y.L., C.Z., and K.

