## Supplementary Materials for "Prognosis prediction using autophagy gene expression in osteosarcoma"

***Data Supplement***

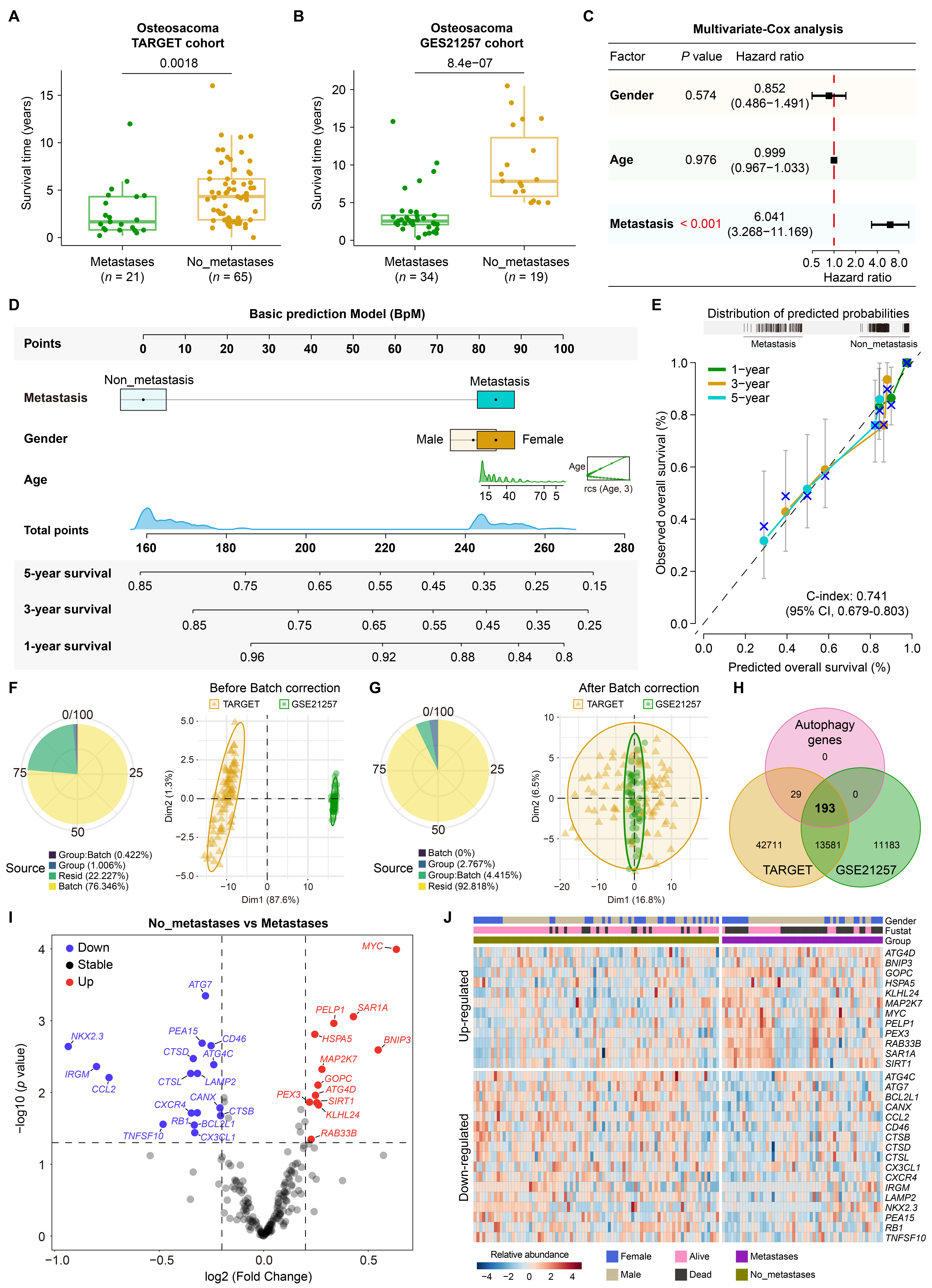

**Fig.S1: Autophagy gene dysregulation in OS**

**A,B**, Survival analyses of the TARGET and GSE21257 cohorts, showing shorter survival time in metastatic patients compared to non-metastatic patients. P values were calculated using Wilcoxon rank-sum test.

**C**, Multivariate COX analyses of OS survival based on gender, age, and metastatic status. Hazard ratio > 1 indicates risk factors, < 1 indicates protective factors. The P values for each variable were derived from the Wald test.

**D**, Overview of the contributions of gender, age, and metastatic status as integrated into the basic prediction model (BpM) and visualized in the nomogram. Each clinical variable was assigned a specific number of points as indicated on the topmost line of the nomogram. The points for all variables were summed to generate a total score, which was then mapped to the probabilities of 1-, 3-, and 5-year overall survival. The knot values displayed in the inset RCS plot correspond to the positions selected for spline fitting; note that they are not arranged sequentially by age along the axis.

**E**, Calibration plots for the validation of the BpM. The predicted probability of overall survival (x-axis, as estimated by the nomogram) is plotted against the observed overall survival (y-axis, based on Kaplan–Meier estimates). Vertical grey lines indicate the 95% confidence intervals. The black dashed diagonal line represents the ideal reference line, where predictions perfectly match observations. Blue “×” marks represent pointwise calibration results, and short vertical ticks at the top denote the distribution of predicted probabilities across individual samples.

**F,G**, Variance decomposition (pie charts) and principal component analysis (PCA) of the TARGET (yellow) and GSE21257(green) cohorts shown before (F) and after (G) batch effect correction. Variance was attributed to Group (metastasis vs non-metastasis), Batch (cohort differences), Group:Batch (interaction), and Resid (unexplained). PCA plots show distinct separation between cohorts before correction and improved integration after batch effect removal.

**H**, Venn diagram illustrating the overlap of autophagy genes, curated from the Human Autophagy Database, and those detected in the TARGET and GSE21257 transcriptional cohorts.

**I**, The changes in autophagy gene expression in the integrated cohorts comparing metastatic and non-metastatic group.

**J**, Abundance trajectories of all 29 changed autophagy genes were normalized by row-wise Z-score across samples and clustered according to metastatic status. The distribution of gender and fustat (survival status) among 139 OS patients is shown horizontally above.

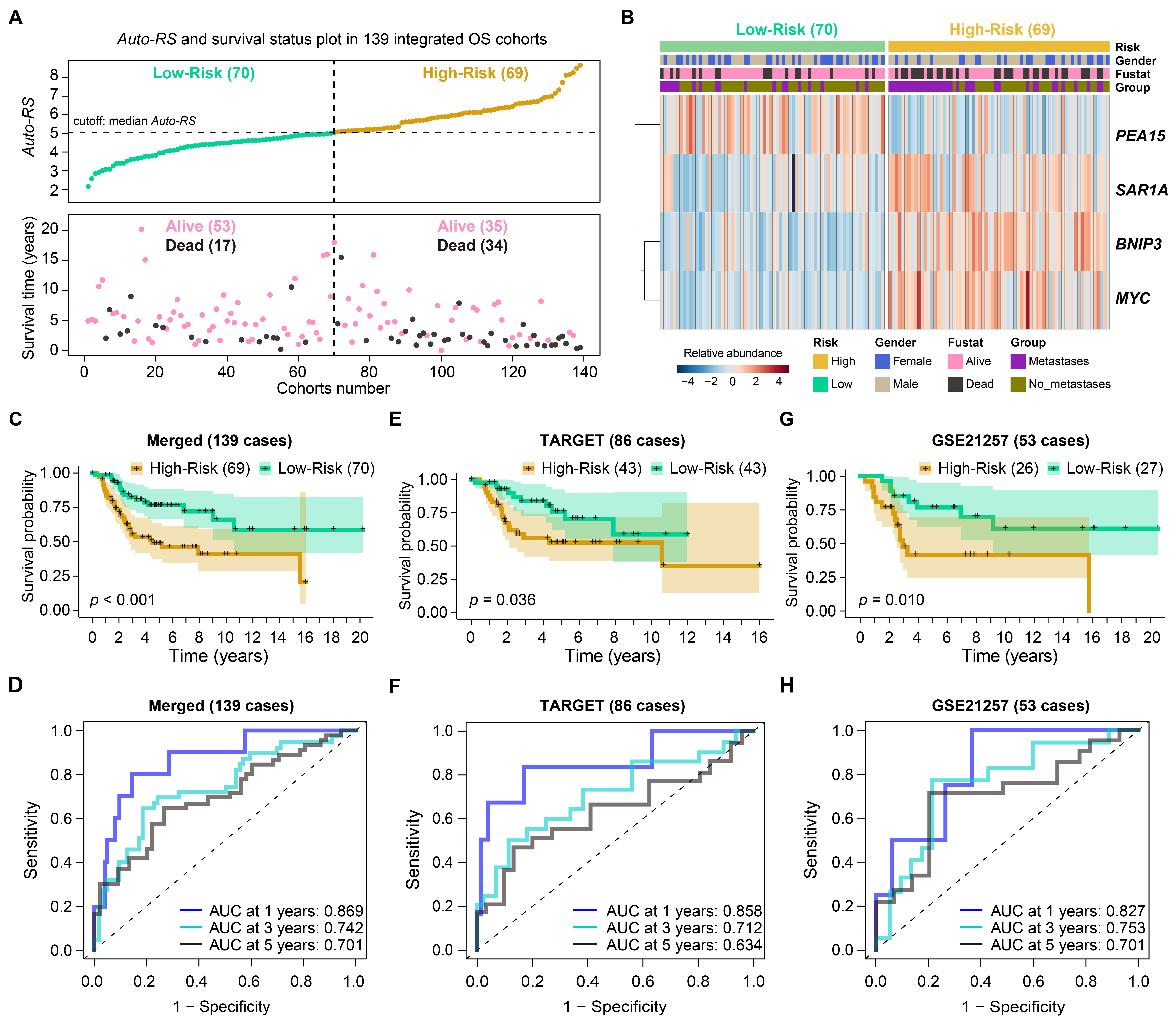

**Fig.S2: Prognostic models integrating** **Auto-RS and clinical variables in OS**

**A**, Univariate Cox regression of OS survival for gender, age, Auto-RS, and metastasis. Horizontal lines indicate the 95% CI of hazard ratios. P values were calculated using a two-sided likelihood ratio test.

**B**, Multivariate Cox regression incorporating gender, age, Auto-RS, and metastasis.

**C**, ROC curves comparing predictive performance of Auto-RS and clinical variables. AUC indicates discrimination; values closer to 1 reflect higher accuracy.

**D**, Nomogram of the complete prediction model (CpM) integrating gender, age, Auto-RS, and metastasis to estimate 1-, 3-, and 5-year OS. Points assigned to each variable (top line) are summed into a total score, then mapped to survival probabilities. Inset RCS plot shows spline knot positions (not sequentially aligned by age).

**E**, Calibration plot of CpM. Predicted OS (x-axis) versus observed OS (y-axis, Kaplan–Meier). Black dashed line, ideal concordance; blue ×, pointwise calibration; vertical ticks, predicted probability distribution.

**F**, Nomogram of the autophagy prediction model (ApM), integrating Auto-RS, age, and gender (excluding metastasis) for 1-, 3-, and 5-year OS estimation. Point assignment and total score mapping as in D.

**G**, Calibration plot for ApM, conventions as in E.

**H**, Concordance index (C-index) comparison among CpM, BpM, and ApM. Higher C-index indicates better discrimination. P values were calculated using a likelihood ratio test.

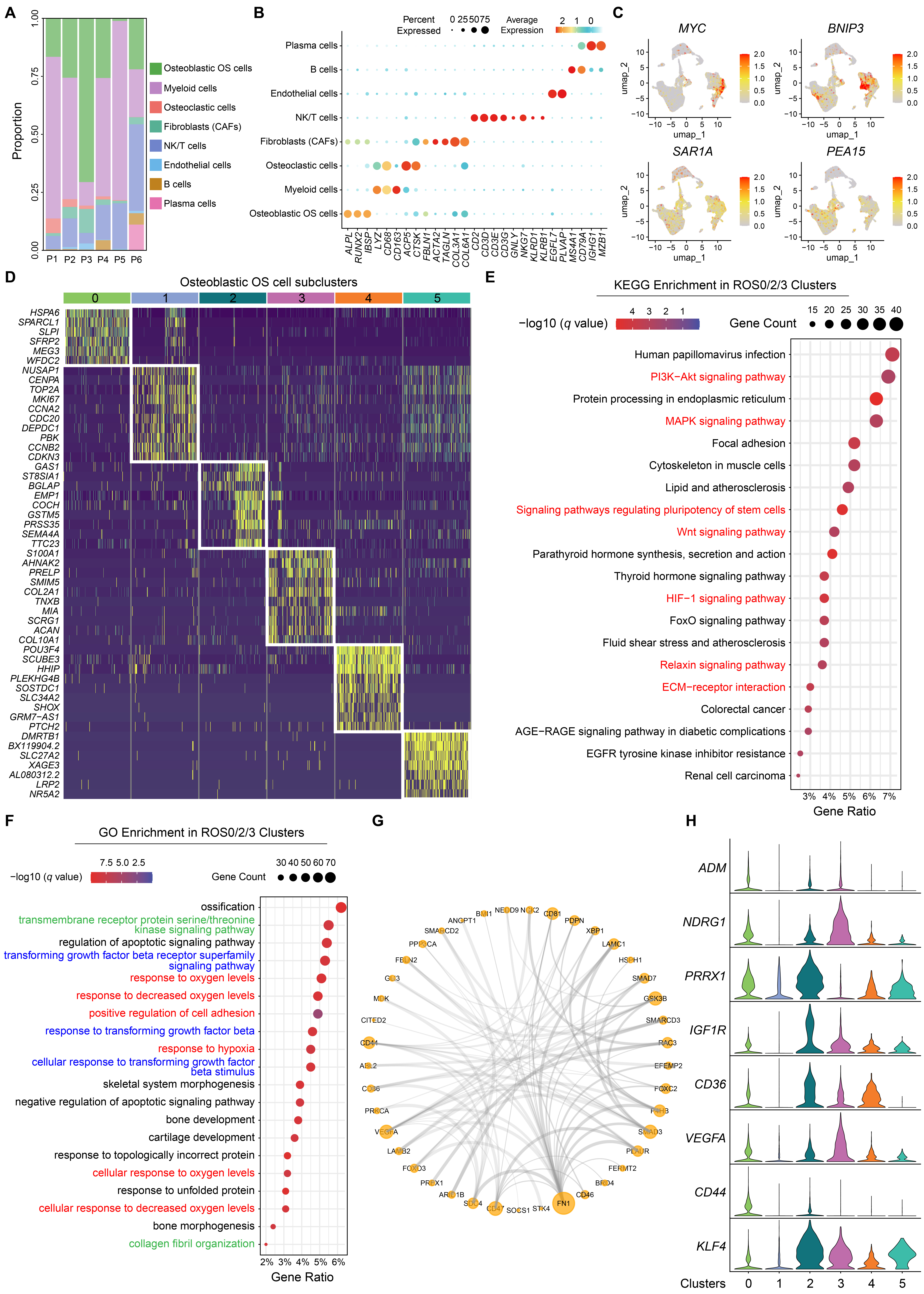

**Fig. S3**: **Comprehensive characterization of osteoblastic OS cell subclusters.**

**A**, Stacked bar chart of OS cluster cell proportions in six patients.

**B**, Dot plot of representative marker genes across eight cell subtypes. Dot size indicates the proportion of expressing cells; color reflects relative expression.

**C**, UMAP visualization of the four model genes.

**D**, Heatmap of the top ten marker genes in the osteoblastic OS subtype.

**E**, KEGG enrichment of genes upregulated in ROS0/2/3 clusters (top 20 pathways). Dot size shows the number of mapped genes; color indicates significance.

**F**, GO enrichment of genes upregulated in ROS0/2/3 clusters (top 20 terms). Dot size shows the number of mapped genes; color indicates significance.

**G**, Protein–protein interaction network of genes in the “positive regulation of cell adhesion” GO term.

**H**, Violin plot of metastasis-associated genes in the osteoblastic OS subtype.

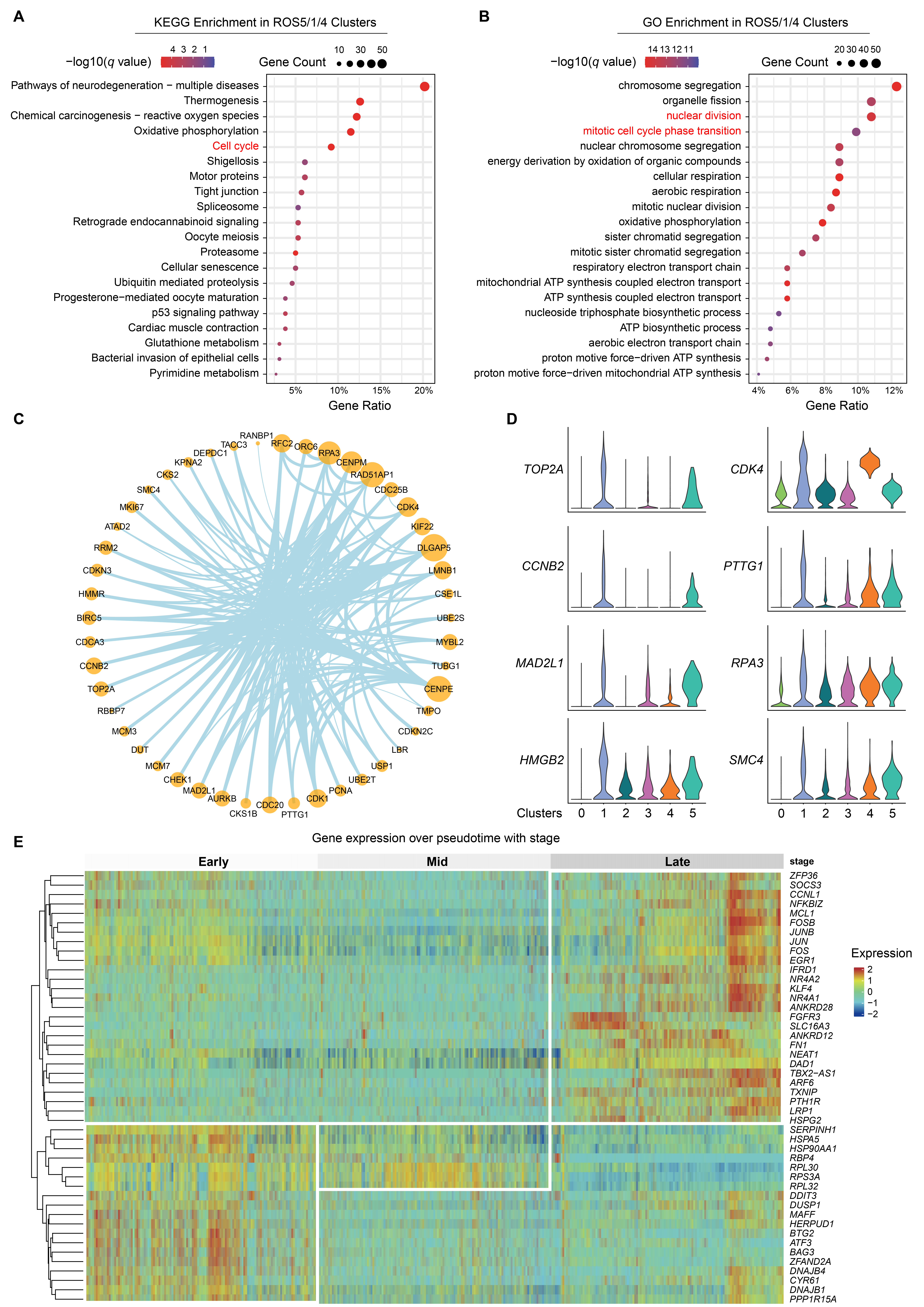

**Fig. S4: Comprehensive analysis of functional enrichment, interaction networks, and pseudotime expression in osteoblastic osteosarcoma**

**A**, KEGG enrichment of genes clusters (top 20 pathways). Dot size shows the number upregulated in ROS5/1/4 of mapped genes; color I ndicates significance.

**B**, GO enrichment of genes upregulated in ROS5/1/4 clusters (top 20 terms). Dot size shows the number of mapped genes; color indicates significance.

**C**, Protein–protein interaction network of genes in the “cell cycle” GO term.

**D**, Violin plot of proliferation-associated genes in the osteoblastic OS subtype.

**E**, Heatmap of pseudotime dynamics of representative differentially expressed genes. Rows are genes, columns are cells; color indicates scaled expression. Top annotation denotes pseudotime stages (Early, Mid, Late).

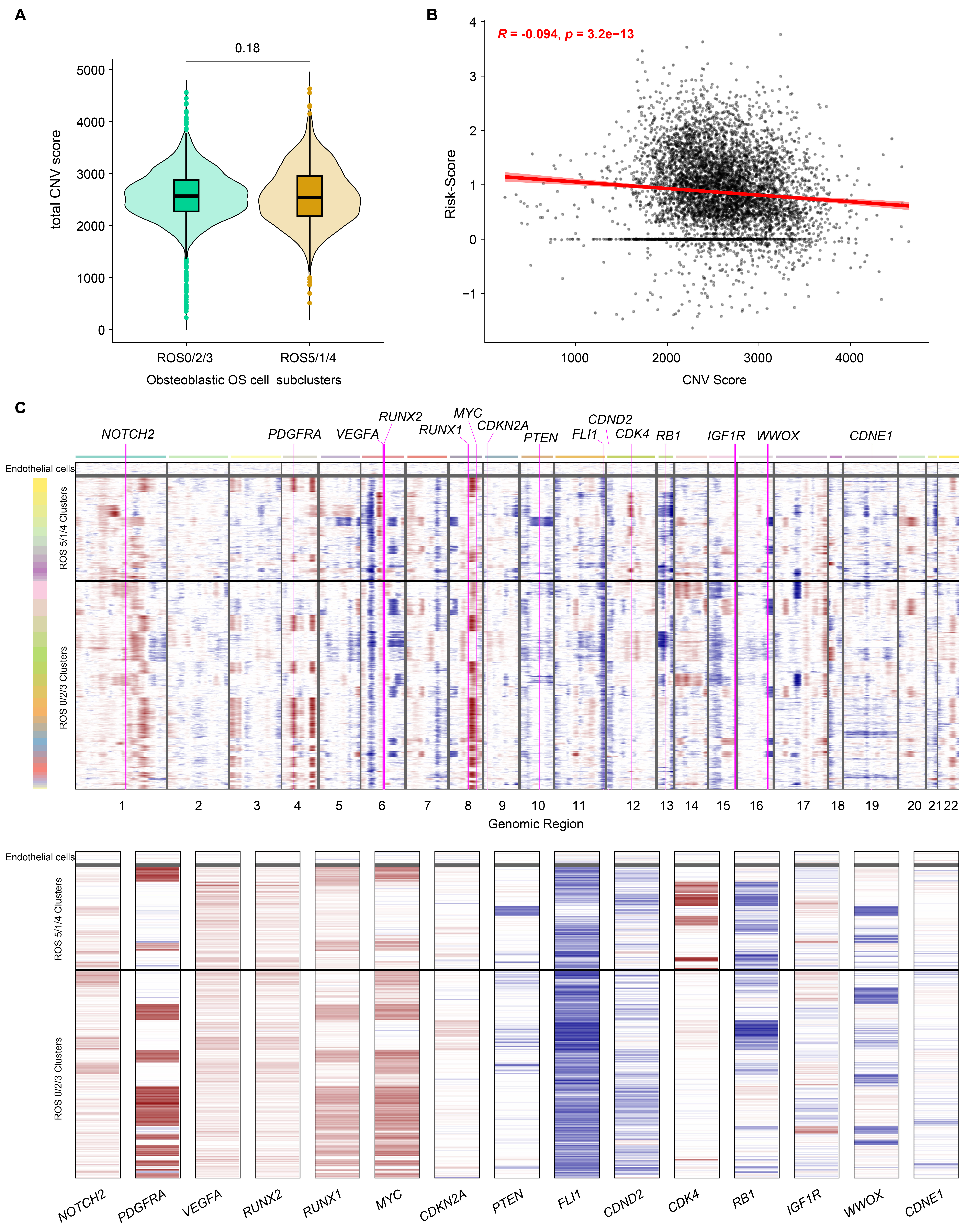

**Fig. S5:** **Relationship between *Auto-RS* and CNV in osteoblastic OS cells**

**A**, Total CNV scores compared between ROS0/2/3 and ROS5/1/4 clusters (Wilcoxon rank-sum test).

**B**, Correlation between CNV scores and *Auto-RS* assessed by Spearman’s rank correlation.

**C**, Next-generation clustered heatmap (NG-CHM). Pink lines mark selected chromosomal segments; the lower panel shows a magnified view of the corresponding region.

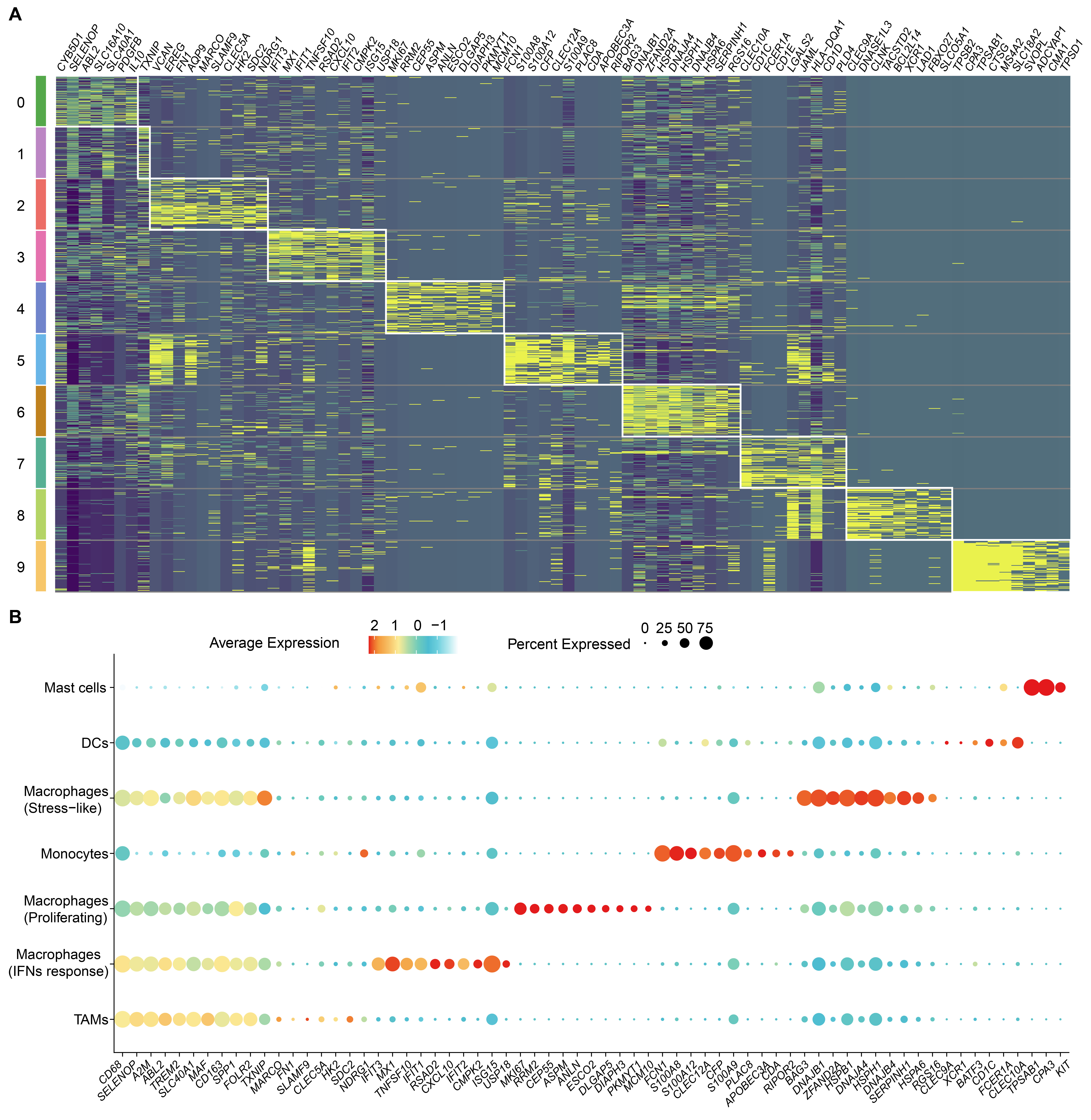

**Fig. S6:** **Marker gene expression patterns of myeloid cell subtypes.**

**A**, Heatmap of the top ten marker genes in myeloid subtypes (RMC0–RMC9).

**B**, Dot plot of representative marker genes across myeloid clusters. Dot size indicates the fraction of expressing cells; color reflects relative expression.

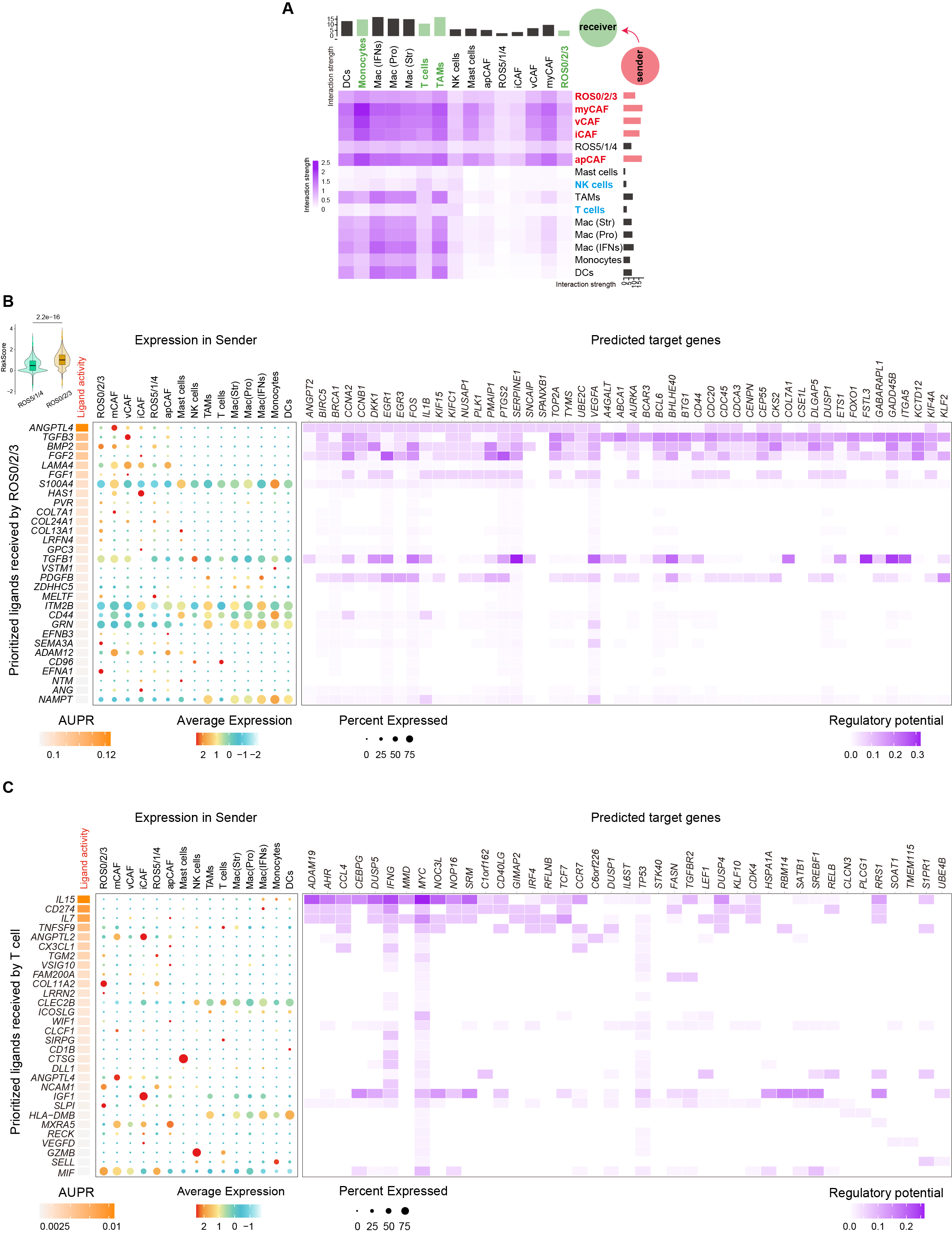

**Fig. S7:** **Cell–cell communication analysis in the osteosarcoma microenvironment.**

**A**, Ligand–receptor interaction strength among osteoblastic OS, CAF, myeloid, NK, and T cell subtypes. Rows represent sender cells (ordered by *Auto-RS*, low to high, bottom to top); columns represent receiver cells (ordered by *Auto-RS*, low to high, left to right).

**B**, NicheNet analysis of ligand–target signaling with ROS0/2/3 as receiver and surrounding clusters as senders. Left: ligands; right: predicted target genes. The violin plot (upper left) shows *Auto-RS* distributions between ROS0/2/3 and ROS5/1/4 clusters, with ROS0/2/3 having significantly higher scores (*P* < 0.001).

**C**, NicheNet analysis of ligand–target signaling with T cells as receiver and surrounding clusters as senders. Left: ligands; right: predicted target genes. The violin plot (upper left) shows *Auto-RS* distributions between low- and high-risk groups, with high-risk groups having significantly higher scores (*P* < 0.001).

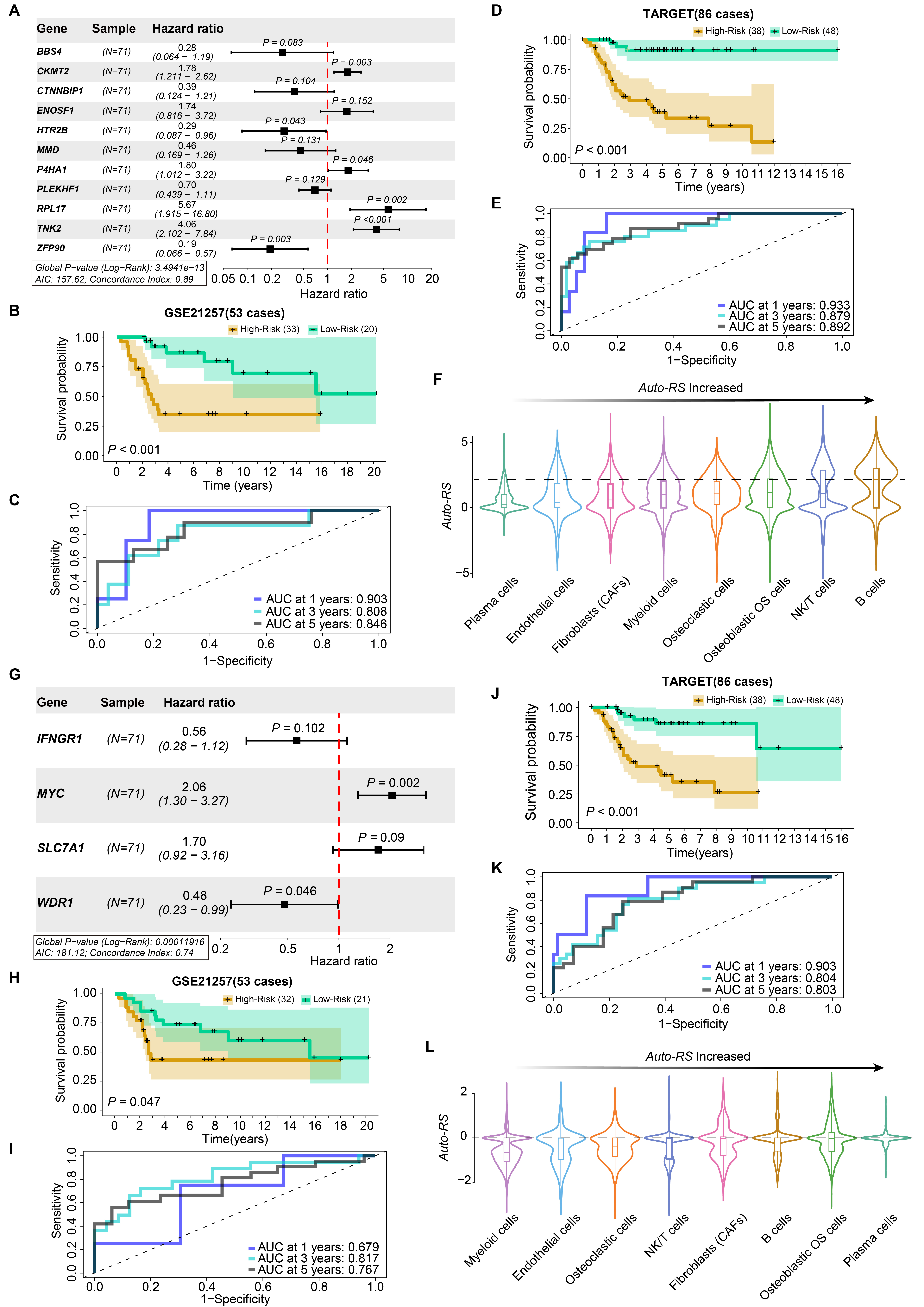

**Fig. S8: Model analysis of genes enriched in the whole transcriptome and tumor-related metastasis pathways**

**A**, Forest plot of 11 whole-transcriptome genes identified by multivariate Cox regression. Horizontal lines show 95% confidence intervals (CI) of hazard ratios. The red dashed line marks hazard ratios = 1 (no effect), > 1 indicates risk, < 1 indicates protection. *P* values were derived from the Wald test.

**B**,**C**, Kaplan–Meier survival curves (**B**) and ROC analysis (**C**) of the predictive model in GSE21257 (*n* = 53). High-risk patients showed significantly shorter survival (log-rank test), with ROC curves indicating good predictive accuracy at 1, 3, and 5 years.

**D**,**E**, Kaplan–Meier survival curves (**D**) and ROC analysis (**E**) in TARGET (*n* = 86), showing consistent results.

**F**, *Auto-RS* based on the 11-gene model applied to single-cell data. Distribution of *Auto-RS* across eight major cell populations, increasing from left to right.

**G**, Forest plot of four metastasis-related genes identified by multivariate Cox regression Horizontal lines show 95% confidence intervals (CI) of hazard ratios. The red dashed line marks hazard ratios = 1 (no effect), > 1 indicates risk, < 1 indicates protection. *P* values were derived from the Wald test.

**H**,**I**, Kaplan–Meier survival curves (**H**) and ROC analysis (**I**) of the 4-gene model in GSE21257 (*n* = 53). High-risk patients showed significantly shorter survival (log-rank test), with ROC curves indicating good predictive accuracy at 1, 3, and 5 years.

**J**,**K**, Kaplan–Meier survival curves (**J**) and ROC analysis (**K**) in TARGET (*n* = 86). High-risk patients showed significantly shorter survival (log-rank test), with ROC curves indicating good predictive accuracy at 1, 3, and 5 years.

**L**, *Auto-RS* based on the 4-gene model applied to single-cell data. *Auto-RS* distribution across eight major cell populations, increasing from left to right.
